# Ursodeoxycholic acid (UDCA) mitigates the host inflammatory response during *Clostridioides difficile* infection by altering gut bile acids which attenuates NF-κB signaling via bile acid activated receptors

**DOI:** 10.1101/2020.01.16.910018

**Authors:** Jenessa A. Winston, Alissa J. Rivera, Jingwei Cai, Rajani Thanissery, Stephanie A. Montgomery, Andrew D. Patterson, Casey M. Theriot

## Abstract

*Clostridioides difficile* infection (CDI) is associated with increasing morbidity and mortality posing an urgent threat to public health. Recurrence of CDI after successful treatment with antibiotics is high, thus necessitating discovery of novel therapeutics against this enteric pathogen. Administration of the secondary bile acid ursodeoxycholic acid (UDCA, ursodiol) inhibits the life cycle of various strains of *C. difficile in vitro*, suggesting the FDA approved formulation of UDCA, known as ursodiol, may be able to restore colonization resistance against *C. difficile in vivo*. However, the mechanism(s) by which ursodiol is able to restore colonization resistance against *C. difficile* remains unknown. Here, we confirmed that ursodiol inhibits *C. difficile* R20291 spore germination and outgrowth, growth, and toxin activity in a dose dependent manner *in vitro*. In a murine model of CDI, exogenous administration of ursodiol resulted in significant alterations in the bile acid metabolome with little to no changes in gut microbial community structure. Ursodiol pretreatment resulted in attenuation of CDI pathogenesis early in the course of disease, which coincided with alterations in the cecal and colonic inflammatory transcriptome, bile acid activated receptors nuclear farnesoid X receptor (FXR), and transmembrane G protein-coupled membrane receptor 5 (TGR5), which are able to modulate the innate immune response through signaling pathways such as NF-κB. Although ursodiol pretreatment did not result in a consistent decrease in the *C. difficile* life cycle *in vivo*, it was able to attenuate an overly robust inflammatory response that is detrimental to the host during CDI. Ursodiol remains a viable non-antibiotic treatment and/or prevention strategy against CDI. Likewise, modulation of the host innate immune response via bile acid activated receptors, FXR and TGR5, represents a new potential treatment strategy for patients with CDI.

**Importance:** The clinical utility of ursodiol for prevention of recurrent CDI is currently in Phase 4 clinical trials. However, the mechanism by which ursodiol exerts its impacts on *C. difficile* pathogenesis is poorly understood. Herein, we demonstrated that ursodiol pretreatment attenuates CDI pathogenesis early in the course of disease in mice, which coincides with alterations in the cecal and colonic inflammatory transcriptome, bile acid activated receptors nuclear farnesoid X receptor (FXR), and transmembrane G protein-coupled membrane receptor 5 (TGR5), which are able to modulate the innate immune response through signaling pathways such as NF-κB. Ursodiol attenuated an overly robust inflammatory response that is detrimental to the host during CDI, and thus remains a viable non-antibiotic treatment and/or prevention strategy against CDI. Likewise, modulation of the host innate immune response via bile acid activated receptors, FXR and TGR5, represents a new potential treatment strategy for patients with CDI.

**Abbreviations:** αMCA – α-Muricholic acid; βMCA –β-Muricholic acid; ωMCA –ω-Muricholic acid; CA – Cholic acid; CDCA – Chenodeoxycholic acid; DCA – Deoxycholic acid; GCDCA – Glycochenodeoxycholic acid; GDCA – Glycodeoxycholic acid; GLCA – Glycolithocholic acid; GUDCA – Glycoursodeoxycholic acid; HCA – Hyodeoxycholic acid; iDCA – Isodeoxycholic acid; iLCA – Isolithocholic acid; LCA – Lithocholic acid; TCA – Taurocholic acid; TCDCA – Taurochenodeoxycholic acid; TDCA – Taurodeoxycholic acid; THCA – Taurohyodeoxycholic acid; TUDCA – Tauroursodeoxycholic acid; TβMCA– Tauro-β-muricholic acid; TωMCA –Tauro ω-muricholic acid; UDCA Ursodeoxycholic acid.

## Introduction

*Clostridioides difficile* is a leading nosocomial enteric pathogen that causes significant human morbidity and mortality, resulting in 29,000 deaths per year and costing over $4.8 billion in healthcare expenses annually in the United States.^1–5^ A major risk factor for *C. difficile* infection (CDI) is the use of antibiotics.^6, 7^ Antibiotics lead to significant and enduring shifts in the gut microbiota and metabolome^8–10^, resulting in loss of colonization resistance against *C. difficile*. Although the exact mechanisms of colonization resistance against *C. difficile* remain unclear, there is increasing evidence that microbial derived secondary bile acids play an important role.^10–13^

Antibiotics, either vancomycin or fidaxomicin, are recommended for an initial episode of CDI.^14^ Unfortunately antibiotic treatment can further disrupt the gut microbial community and recurrence of CDI after cessation of antibiotics is high, occurring in 20-30% of patients.^3, 15–18^ Even fidaxomicin, a bactericidal antibiotic against *C. difficile,* alters the intestinal microbial community structure, just to a lesser extent than vancomycin.^19^ Fecal microbiota transplantation (FMT) is a highly effective treatment for recurrent CDI, however the long-term ramifications of this conditionally FDA approved procedure are unknown.^20, 21^ A potential mechanism by which FMT leads to resolution of CDI in a mouse model^22, 23^ and in humans^13^ is by restoring microbial derived secondary bile acids.

Secondary bile acids, such as deoxycholic acid (DCA), lithocholic acid (LCA) and ursodeoxycholic acid (UDCA, ursodiol), are produced as a collaborative effort by the host (production of primary bile acids) and the gut microbiota (biochemical modification of host primary bile acids into secondary bile acids).^24, 25^ Our group and others have demonstrated that numerous microbial derived secondary bile acids, including UDCA, are able to inhibit spore germination, growth, and toxin activity of various *C. difficile* strains *in vitro*.^26, 27^ We also know that secondary bile acids are associated with colonization resistance against *C. difficile* in mice and humans.^10, 13, 28^ Furthermore, oral UDCA administration successfully prevented recurrence of *C. difficile* ileal pouchitis in a single patient,^29^ suggesting that exogenous administration of the secondary bile acid UDCA may be able to restore colonization resistance against *C. difficile in vivo*.^29^

UDCA is available in an FDA approved formulation known as ursodiol.^30^ Ursodiol is used to treat a variety of hepatic and biliary diseases due to its myriad of beneficial effects, including: anticholestatic, antifibrotic, antiproliferative, and anti-inflammatory properties.^31–40^ Additionally, off-label use of ursodiol is currently in Phase 4 clinical trials for the prevention of recurrent CDI.^41^ It is well established that ursodiol alters the hepatic, biliary, and serum bile acid pools.^31, 42^ However, it remains unclear if and how exogenously administered ursodiol alters the indigenous gut microbial community structure and/or bile acid metabolome, which contributes to colonization resistance against *C. difficile*.

Based on our previous work with UDCA,^26^ we hypothesized that ursodiol would *directly* inhibit different stages of the *C. difficile* life cycl*e,* and ameliorate disease when administered to mice in a model of CDI. Due to their detergent like properties, bile acids can also directly alter the gut microbial composition, subsequently altering bile acid metabolism and host physiology.^43^ Gastrointestinal crosstalk between bile acids and the gut microbiota is well recognized (reviewed in Wahlström et al.)^43^, but the impact of ursodiol on these interactions remains unknown. Similarly, the central role bile acids play in regulation of the innate immune response is growing (reviewed in Fiorucci et al.).^44^ Because of this, we also investigated the *indirect* effects or the extent to which the impact of ursodiol on CDI correlated with changes in the gut microbiota, bile acid metabolome, and the host inflammatory response.

Here we show that ursodiol is able to significantly decrease *C. difficile* spore germination, growth, and toxin activity in a dose-dependent manner *in vitro,* but not to the same level *in vivo*. In *C. difficile* infected mice, ursodiol pretreatment resulted in attenuation of CDI pathogenesis early in the course of disease, which coincided with alterations in gut bile acids, the host inflammatory transcriptome, and gene expression of bile acid activated receptors nuclear farnesoid X receptor (FXR) and transmembrane G protein-coupled membrane receptor 5 (TGR5), which are able to modulate the innate immune response through signaling pathways such as NF-κB. Although ursodiol pretreatment did not restore colonization resistance against *C. difficile*, it attenuated an overly robust host inflammatory response that is detrimental to the host during CDI. Together, these results suggest that ursodiol remains a viable non-antibiotic treatment and/or prevention strategy against CDI by acting on the FXR-FGR axis via NF-kB signaling. Targeting the bile acid activated receptors, FXR and TGR5, represents a novel approach for the development of therapies for the prevention and treatment of CDI.

## Materials and Methods

### Spore preparation

*C. difficile* R20291 spores were prepared as described previously.^22, 26, 45^ Briefly *C. difficile* R20291 were grown at 37°C anaerobically overnight in 2 ml Columbia broth and then added to 40 ml culture of Clospore media in which it was allowed to sporulate for 5–7 days. Spores were harvested by centrifugation, and subjected to 3–5 washes with sterile cold water to ensure spore purity. Spore stocks were stored at 4°C in sterile water until use.

### *C. difficile in vitro* spore germination and outgrowth assay

The spore germination and outgrowth assay performed was modified from Carlson et al., Theriot et al., and Thanissery et al.^22, 26, 46^ Purified spores were enumerated and tested for purity before use. The spore stock was resuspended in ultrapure water to achieve an initial spore concentration of approximately 10^6^ spores/ml. The spore suspension was subjected to heat treatment (65°C for 20 min) to eliminate any vegetative cells. The spores were then plated on both brain heart infusion (BHI) with 100 mg/liter L-cysteine, and BHI media supplemented with 0.1% TCA, and further incubated at 37°C for 24 hr. We observed a lawn of *C. difficile* colonies on BHI media supplemented with 0.1% TCA, and no visible growth on BHI media alone, which indicates that the spore suspension was devoid of vegetative cells. Bile acid solutions containing four to five concentrations were used to study a dose response [Ursodiol U.S.P (0.0076, 0.076, 0.76, and 7.64 mM, Spectrum Chemical, CAS 128-13-2)]. Bile acids were dissolved in ethanol, passed into the anaerobic chamber (Coy labs), and added to BHI broth with 0.1% TCA. Chendeoxycholate (CDCA, Fisher Scientific, 50328656) at 0.04% is a known inhibitor of TCA mediated spore germination, and was used as a negative control. TCA at 0.1% made in water, and water and ethanol were used as positive controls. *C. difficile* R20291 spores were added to BHI broth supplemented with and without TCA 0.1% and bile acids at concentrations listed above, and allowed to incubate for 30 min at 37°C anaerobically. Bacterial enumeration of the samples was performed on both BHI agar (vegetative cells only) and BHI agar supplemented with 0.1% TCA (germinated spores and vegetative cells). Percent germination was calculated as [(CFUs on BHI supplemented with TCA + SBAs) / (CFUs on BHI supplemented with TCA alone)] × 100. All measurements were performed in triplicate for each isolate and expressed as percent germination.

### *C. difficile in vitro* growth kinetics assay

The growth kinetic assay was modified from Thanissery et al. and performed over a 24 hr period.^26^ *C. difficile* R20291 was cultured overnight at 37°C in BHI plus 100 mg/liter L-cysteine broth in an anaerobic chamber. After 14 hr of growth, *C. difficile* R20291 was subcultured 1:10 and 1:5 into BHI plus 100 mg/liter L-cysteine and allowed to grow for 3 hr anaerobically at 37°C. The culture was then diluted in fresh BHI to a starting optical density at 600 nm (OD_600_) of 0.01 within a conical tube at a final volume of 50 ml. Five concentrations of filter-sterilized ursodiol (0.0076, 0.076, 0.76, 3.82, 7.64 mM dissolved in ethanol) were added to each culture and anaerobically incubated for 72 hr at 37°C. The optical density was monitored every 4 hours for 12 hr and then once at 24 hr. Cultures were inverted prior to obtaining the optical density on a cell density meter (WPA Biowave). Due to optical density interference from gelatin formation at 7.64 mM ursodiol, this concentration was only assessed for viability.

To assess *C. difficile* R20291 viability, culture aliquots (100 μL) were taken at each time point and enumerated on TBHI plates to obtain total colony forming units (CFU)/mL, which represents total vegetative cells and spores of *C. difficile* R20291. At each time point, a second culture aliquot (100 μL) was heat-treated at 65°C for 20 min in order to eliminate all vegetative cells. This heat-treated culture aliquot was then enumerated on TBHI plates to obtain total CFU/ml, which represents total spores only.

### *C. difficile in vitro* Vero cell cytotoxicity assay

This protocol is modified from Winston et al.^47^ Vero cells were grown and maintained in DMEM media (Gibco Laboratories, 11965-092) with 10% fetal bovine serum (Gibco Laboratories, 16140-071) and 1% Penicillin streptomycin solution (Gibco Laboratories, 15070-063). Cells were incubated with 0.25% trypsin (Gibco Laboratories, 25200-056) washed with 1X DMEM media and harvested by centrifugation 1,000 RPM for 5 min. Cells were plated at 1 × 10^4^ cells per well in a 96-well flat bottom microtiter plate (Corning, 3596) and incubated overnight at 37°C / 5% CO_2_. Culture supernatants from the anaerobic growth assay were defrosted on ice and three 200 μL aliquots from each treatment and time point were then centrifuged at 1,750 RPM for 5 min to pellet vegetative *C. difficile*. Supernatants were collected from each sample and 10-fold dilutions, to a maximum of 10^−6^, were performed. Sample dilutions were incubated 1:1 with PBS (for all dilutions) or antitoxin (performed for 10^−1^ and 10^−4^ dilutions only, TechLabs, T5000) for 40 min at room temperature. Following incubation, these admixtures were added to the Vero cells and plates were incubated overnight at 37°C / 5% CO_2_. Vero cells were viewed under 200X magnification for rounding after overnight incubation. The cytotoxic titer was defined as the reciprocal of the highest dilution that produced rounding in 80% of Vero cells for each sample. Vero cells treated with purified *C. difficile* toxins (A and B) and antitoxin (List Biological Labs, 152C and 155C; TechLabs, T5000) were used as controls.

### Ethics statement

The Institutional Animal Care and Use Committee (IACUC) at North Carolina State University College of Veterinary Medicine (NCSU) approved this study. The NCSU Animal Care and Use policy applies standards and guidelines set forth in the Animal Welfare Act and Health Research Extension Act of 1985. Laboratory animal facilities at NCSU adhere to guidelines set forth in the Guide for the Care and Use of Laboratory Animals. The animals’ health statuses were assessed daily, and moribund animals were humanely euthanized by CO_2_ asphyxiation followed by secondary measures (cervical dislocation). Trained animal technicians or a veterinarian performed animal husbandry in an AAALAC-accredited facility during this study.

### Animals and housing

C57BL/6J mice (females and males) were purchased from Jackson Laboratories (Bar Harbor, ME) and quarantined for 1 week prior to starting the antibiotic water administration to adapt to the new facilities and avoid stress-associated responses. Following quarantine, the mice were housed with autoclaved food, bedding, and water. Cage changes were performed weekly by laboratory staff in a laminar flow hood. Mice had a 12 hr cycle of light and darkness.

### Ursodiol dosing experiment and sample collection

Groups of 5 week old C57BL/6J mice (male and female) were treated with ursodiol at three doses (50, 150, and 450 mg/kg dissolved in corn oil; Ursodiol U.S.P., Spectrum Chemical, CAS 128-13-2) given daily via oral gavage for 14 days. Concurrently mice were administered cefoperazone (0.5 mg/ml), a broad spectrum antibiotic, in their drinking *ad libitum* for 5 days. A control group of mice were only administered cefoperazone in their drinking water *ad libitum* for 5 days. Two independent experiments were performed, with a total of n = 8 mice/treatment group. Fecal pellets were collected twice daily, flash-frozen and stored at −80°C until further analysis.

### Murine model of *C. difficile* infection and sample collection

Groups of 5 week old C57BL/6J mice (male and female, n = 12 mice/treatment group) were given cefoperazone (0.5 mg/ml), in their drinking *ad libitum* for 5 days followed by a 2-day wash out with regular drinking water *ad libitum* (Fig. 3A). A group of cefoperazone treated mice were pretreated with ursodiol (450 mg/kg dissolved in corn oil; Ursodiol U.S.P., Spectrum Chemical, CAS 128-13-2) daily via oral gavage. Mice were then challenged with 10^5^ spores of *C. difficile* R20291 via oral gavage at Day 0 (*C. diff* + U450). Mice were monitored for weight loss and clinical signs of CDI (lethargy, inappetence, diarrhea, and hunched posture) for 7 days post challenge. Fecal pellets were collected daily for *C. difficile* enumeration. Controls for this experiment included cefoperazone treated mice, no ursodiol, challenged with *C. diff* (*C. diff* only), and cefoperazone treated mice pretreated with the corn oil vehicle (equivalent volume, 75 μl) daily via oral gavage and then challenged with 10^5^ spores of *C. difficile* R20291 via oral gavage at Day 0 (*C. diff +* corn oil). Uninfected controls included cefoperazone treated mice with (Cef only), and with pretreatment of ursodiol (450 mg/kg dissolved in corn oil; Cef + U450). Animals were euthanized after losing 20% of initial baseline weight or after developing any serve clinical signs noted above.

Necropsy was performed on days 2, 4, and 7 post challenge. Contents and tissue from the cecum and colon were collected immediately at necropsy, flash frozen in liquid nitrogen, and stored at −80°C until further analysis. To prevent RNA degradation, snips of cecal and colonic tissue were also stored in RNA-Later at 80°C until further analysis. Cecum and colon were prepared for histology by placing intact tissue into histology cassettes and stored in 10% neutral buffered formalin at a ratio of 1:10 tissue:fixative at room temperature for 72 hr then transferred to room temperature 70% ethyl alcohol. Tissues were processed, paraffin embedded, sectioned at 4 µm thick, and stained by routine hematoxlyin and eosin for histopathological examination (LCCC Animal Histopathology Core Facility at the University of North Carolina at Chapel Hill). Histolopathologic evaluation was conducted by a board-certified veterinary pathologist (S. Montgomery) in an *a priori* blinded manner in which individuals within groups were masked at time of scoring. For tissue scoring, a 0-4 numerical scoring system was employed to separately assess edema, inflammatory cell infiltration, and epithelial cell damage in the cecum and colon based upon a previously published scoring scheme.^48^

### Bile acid quantitation by UPLC-MS/MS of murine cecal and fecal content

Targeted analysis of bile acids on fecal pellets (n = 3 mice/treatment group were selected) were performed with an ACQUITY ultraperformance liquid-chromatography system using a C8 BEH column (2.1 × 100 mm, 1.7 µm) coupled with a Xevo TQ-S triplequadrupole mass spectrometer equipped with an electrospray ionization (ESI) source operating in negative ionization mode (All Waters, Milford, MA) as previously described.^49^ The sample was thawed on ice and 25 mg was added to 1 mL of pre-cooled methanol containing 0.5 μM stable-isotope-labeled bile acids as internal standards (IS), followed by homogenization (Precellys, Bertin Technologies, Rockville, MD) with 1.0 mm diameter zirconia/silica beads (BioSpec, Bartlesville, OK) and centrifugation (Eppendorf, Hamburg, Germany) at 13 200g, 4 °C, and 20 min. 200 µl of the supernatant was transferred to an autosampler vial. Following centrifugation, the supernatant was transferred to an autosampler vial for quantitation. Bile acids were detected by either multiple reaction monitoring (MRM) (for conjugated bile acid) or selected ion monitoring (SIM) (for non-conjugated bile acid). MS methods were developed by infusing individual bile acid standards. Calibration curves were used to quantify the biological concentration of bile acids. Bile acid quantitation was performed at Penn State University.

Heatmaps and box and whisker plots of bile acid concentrations, and nonmetric multidimensional scaling (NMDS) depicting the dissimilarity indices via Horn distances between bile acid profiles were generated using R packages (http://www.R-project.org).

### Illumina MiSeq sequencing of bacterial communities

Microbial DNA was extracted from murine feces (n = 3 mice/treatment group were selected) using the PowerSoil-htp 96-well soil DNA isolation kit (Mo Bio Laboratories, Inc.). The V4 region of the 16S rRNA gene was amplified from each sample using a dual-indexing sequencing strategy.^50^ Each 20 µl PCR mixture contained 2 µl of 10× Accuprime PCR buffer II (Life Technologies), 0.15 µl of Accuprime high-fidelity Taq (catalog no. 12346094) high-fidelity DNA polymerase (Life Technologies), 2 µl of a 4.0 µM primer set, 1 µl DNA, and 11.85 µl sterile double-distilled water (ddH_2_O) (free of DNA, RNase, and DNase contamination). The template DNA concentration was 1 to 10 ng/µl for a high bacterial DNA/host DNA ratio. PCR was performed under the following conditions: 2 min at 95°C, followed by 30 cycles of 95°C for 20 sec, 55°C for 15 sec, and 72°C for 5 min, followed by 72°C for 10 min. Each 20 µl PCR mixture contained 2 µl of 10× Accuprime PCR buffer II (Life Technologies), 0.15 µl of Accuprime high-fidelity Taq (catalog no. 12346094) high-fidelity DNA polymerase (Life Technologies), 2 µl of 4.0 µM primer set, 1 µl DNA, and 11.85 µl sterile ddH_2_O (free of DNA, RNase, and DNase contamination). The template DNA concentration was 1 to 10 ng/µl for a high bacterial DNA/host DNA ratio. PCR was performed under the following conditions: 2 min at 95°C, followed by 20 cycles of 95°C for 20 sec, 60°C for 15 sec, and 72°C for 5 min (with a 0.3°C increase of the 60°C annealing temperature each cycle), followed by 20 cycles of 95°C for 20 sec, 55°C for 15 sec, and 72°C for 5 min, followed by 72°C for 10 min. Libraries were normalized using a Life Technologies SequalPrep normalization plate kit (catalog no. A10510-01) following the manufacturer’s protocol. The concentration of the pooled samples was determined using the Kapa Biosystems library quantification kit for Illumina platforms (KapaBiosystems KK4854). The sizes of the amplicons in the library were determined using the Agilent Bioanalyzer high-sensitivity DNA analysis kit (catalog no. 5067-4626). The final library consisted of equal molar amounts from each of the plates, normalized to the pooled plate at the lowest concentration.

Sequencing was done on the Illumina MiSeq platform, using a MiSeq reagent kit V2 with 500 cycles (catalog no. MS-102-2003) according to the manufacturer’s instructions, with modifications.^50^ Libraries were prepared according to Illumina’s protocol for preparing libraries for sequencing on the MiSeq (part 15039740 Rev. D) for 2 or 4 nM libraries. The final load concentration was 4 pM (but it can be up to 8 pM) with a 10% PhiX spike to add diversity. Sequencing reagents were prepared according to Illumina’s protocol for 16S sequencing with the Illumina MiSeq personal sequencer.^50^ (Updated versions of this protocol can be found at http://www.mothur.org/wiki/MiSeq_SOP.) Custom read 1, read 2, and index primers were added to the reagent cartridge, and FASTQ files were generated for paired-end reads.

### Microbiome analysis

Analysis of the V4 region of the 16S rRNA gene was done using mothur (version 1.40.1).^50, 51^ Briefly, the standard operating procedure (SOP) at http://www.mothur.org/wiki/MiSeq_SOP was followed to process the MiSeq data. The paired-end reads were assembled into contigs and then aligned to the SILVA 16S rRNA sequence database (release 132)^52, 53^ and were classified to the mothur-adapted RDP training set v16^54^ using the Wang method and an 80% bootstrap minimum to the family taxonomic level. All samples with <500 sequences were removed. Chimeric sequences were removed using UCHIME.^55^ Sequences were clustered into operational taxonomic units (OTU) using a 3% species-level definition. The OTU data were then filtered to include only those OTU that made up 1% or more of the total sequences. The percentage of relative abundance of bacterial phyla and family members in each sample was calculated. A cutoff of 0.03 (97%) was used to define operational taxonomic units (OTU) and Yue and Clayton dissimilarity metric (θYC) was utilized to assess beta diversity. Standard packages in R were used to create NMDS ordination on serial fecal samples.

### mRNA isolation and gene expression analysis

RNA extraction was performed on cecal and colonic tissue using Zymo Research RNA extraction kit (Irvine, CA). After RNA extraction, samples were quantified using an Invitrogen Qubit (Thermo Fisher Scientific) and run on an Agilent Bioanalyzer NanoChip (Santa Clara, CA) to assess quality of RNA. RNA extraction, quantification, and purity assessment were performed at the Microbiome Shared Resource Lab, Duke University, Durham, NC.

The nCounter Mouse Inflammation v2 gene expression panel and custom code set was purchased from NanoString Technologies (Seattle, WA) and consists of 254 inflammation-related mouse genes including six internal reference genes (see Table S1) and 10 custom genes of interest (see Table S2). The nCounter assay was performed using 100 ng of total RNA. Hybridization reactions were performed according to the manufacturer’s instructions with 5 µl diluted sample preparation reaction and samples were hybridized overnight. Hybridized reactions were purified using the nCounter Prep Station (NanoString Technologies) and data collection was performed on the nCounter Digital Analyzer (NanoString Technologies) following the manufacturer’s instructions to count targets. Nanostring nCounter Max platform was utilized at UNC Translational Genomics Laboratory to perform these assays.

The raw data was normalized to six internal reference genes within each tissue type with the lowest coefficient of variation using nSolver software following the manufacturer’s instructions (Nanostring Technologies). Using the nSolver Advanced analysis software, after pre-processing and normalization counts were log_2_ transformed and z-scores were calculated (Nanostring Technologies).

To identify highly significant genes and molecular pathways, the gene set was subsequently uploaded to Ingenuity Pathway Analysis (IPA) database (Qiagen, Redwood City, CA) and analyzed in the context of known biological response and regulatory networks. IPA was also used to assess gene expression fold changes comparing ursodiol pretreated CDI mice to untreated CDI mice.

### Statistical analysis

Statistical tests were performed using Prism version 7.0b for Mac OS X (GraphPad Software, La Jolla California USA) or using R packages (http://www.R-project.org). Statistical significance was set at a p value of < 0.05 for all analyses (*, p <0.05; **, p < 0.01; ***, p < 0.001; ****, p < 0.0001). For *in vitro* experiments, significance between treatments in spore germination was calculated by Student’s parametric t-test with Welch’s correction. Two-way analysis of variance (ANOVA) test followed by Dunnett’s multiple comparisons *post hoc* test was used to calculate significance between treatments for the anaerobic growth assay and toxin activity assay.

For the *in vivo* experiments, a two-way ANOVA followed by Tukey’s multiple comparisons post hoc test was used to calculate significance between treatment groups in bile acid profiles, individual bile acids, *C. difficile* load (CFU/g) cecal content, and total histologic score. A two-way ANOVA followed by Sidak’s multiple comparisons *post hoc* test was used to calculate significance between treatment groups in baseline weight loss. A Student’s t-test corrected for multiple comparisons using the Holm-Sidak method was used to calculate significance between treatment groups in cecal content toxin activity and edema score. Analysis of molecular variance (AMOVA) was used to detect significant microbial community clustering of treatment groups in NMDS plots.^56^ nSolver software and IPA were used to calculate significance in gene expression fold change comparing mice pretreated with ursodiol to untreated mice.

### Availability of data and material

Raw 16S sequences have been deposited in the Sequence Read Archive (SRA) with SRA accession number: SUB6809988. All other data including bile acid metabolomics and Nanostrings expression data is provided in Tables S3-S7 in the supplemental material.

## Results

### Ursodiol directly inhibits different stages of the *C. difficile* life cycle *in vitro*

To determine the direct effects of ursodiol on the life cycle of clinically relevant *C. difficile* strain R20291, *in vitro* assays to assess spore germination and outgrowth, growth, and toxin activity were performed (Fig. 1). Taurocholic acid (TCA) mediated spore germination and outgrowth of *C. difficile* was significantly inhibited by ursodiol in a dose dependent manner (Fig. 1A, blue bars). No significant difference was observed between positive controls, sterile ethanol, and water supplemented with 0.1% TCA (Fig. 1A, ethanol represented as solid grey bar). As expected, the negative control, 0.04% chenodeoxycholic acid (CDCA) supplemented with 0.1% TCA resulted in significant inhibition of spore germination and outgrowth (Fig. 1A, red bar). CDCA is a known inhibitor of spore germination.^57^

**Figure 1:**
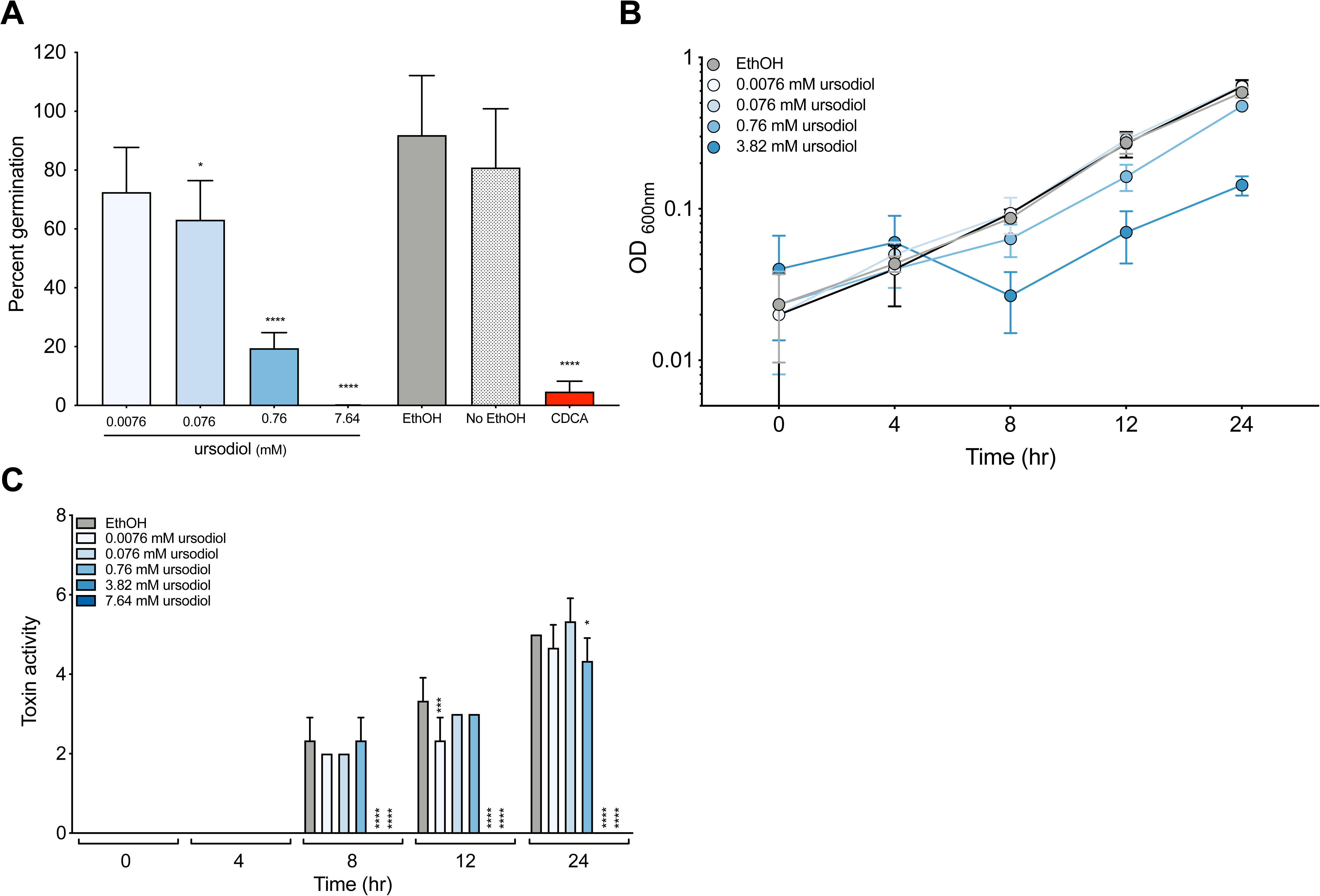
Ursodiol inhibits different stages of the *C. difficile* life cycle. *In vitro* assay to assess if the addition of ursodiol (0.0076, 0.076, 0.76, and 7.64 mM) **(A)** inhibits *C. difficile* R20291 TCA-mediated spore germination and outgrowth compared to positive control, ethanol (gray bar) and negative control, 0.04% CDCA (red bar). The data presented represents triplicate experiments. Statistical significance between treatment groups and the positive controls was determined by Student’s parametric t-test with Welch’s correction. **(B)** alters the growth kinetics of *C. difficile* over a 24 hr period. Growth curves were done in BHI media with ethanol (gray line) and addition of varying concentrations of ursodiol (blue lines). The data presented represents OD_600_ ± SEM from triplicate experiments. **(C)** Culture supernatants, taken throughout a 24 hr growth curve in 1B, were used for a Vero cell cytotoxicity assay and the data is expressed as log_10_ reciprocal dilution toxin per 100 μL of *C. difficile* culture supernatant. The data presented represents triplicate experiments. Statistical significance between treatment groups and the positive controls was determined by a two-way ANOVA with Dunnett’s multiple comparisons (*, p < 0.05; ***, p < 0.001; ****, p < 0.0001).

*C. difficile* growth decreased when exposed to ursodiol at 8, 12 and 24 hr in a dose dependent manner compared to the ethanol control (Fig. 1B, blue lines). For the ursodiol 3.82 mM concentration, a significantly lower optical density (OD) compared to the ethanol control was observed at time points 8, 12, and 24 hr (p = 0.0316, p = 0.0001, p = 0.0001, respectively). For the ursodiol 0.76 mM treatment, a significantly lower OD compared to ethanol control was observed at time points 12 and 24 hr (p = 0.0001 for both time points), and for the ursodiol 0.076 mM treatment at time point 24 hr (p = 0.0208). No significant difference in OD between the ethanol control and treatments was noted at time point 0 or 4 hr. The highest concentration of ursodiol (7.64 mM) interfered with OD measurement and was not included in the analysis.

Since OD measurements do not capture viability and spore formation of *C. difficile*, we also did differential plating to evaluate the colony forming units (CFU)/ml at each time point (Fig. S1). At 24 hr ursodiol inhibited *C. difficile* total vegetative cells and spores compared to the ethanol control. The highest concentration of ursodiol (7.64 mM) had a significant and immediate inhibitory effect on *C. difficile* viability compared to ethanol at time 0 hr. Ursodiol treatment also significantly decreased *C. difficile* spores.

Toxin activity was not detected until eight hours of *C. difficile* growth (Fig. 1C). Higher concentrations of ursodiol resulted in significant decreases in *C. difficile* toxin activity compared to the ethanol control at 8, 12, and 24 hr (Fig. 1C, blue bars). Using a Spearman’s correlation (r) the CFU/ml and toxin activity at 8 and 12 hr were significantly correlated (8 hr, r = 0.6399, p = 0.0042; 12 hr, r = 0.6981, p = 0.0013), suggesting low toxin activity is due to decreased growth.

### Ursodiol alters the fecal bile acid metabolome with minimal change to the microbiota

Since the gut microbiota and bile acids mediate colonization resistance against *C. difficile*, we aimed to determine how different doses of ursodiol administration alone impacts them in our antibiotic treated mouse model. C57BL/6J mice were administered three doses of ursodiol (50, 150, 450 mg/kg/day) via oral gavage for 14 days (Fig. 2A). Mice were also concurrently administered cefoperazone (in drinking water *ad libitum*) for 5 days, in order to mimic the CDI mouse model. Paired fecal samples were collected from the same mice serially over the 14 day period to evaluate how ursodiol affected the gut microbial community structure and bile acid metabolome.

**Figure 2:**
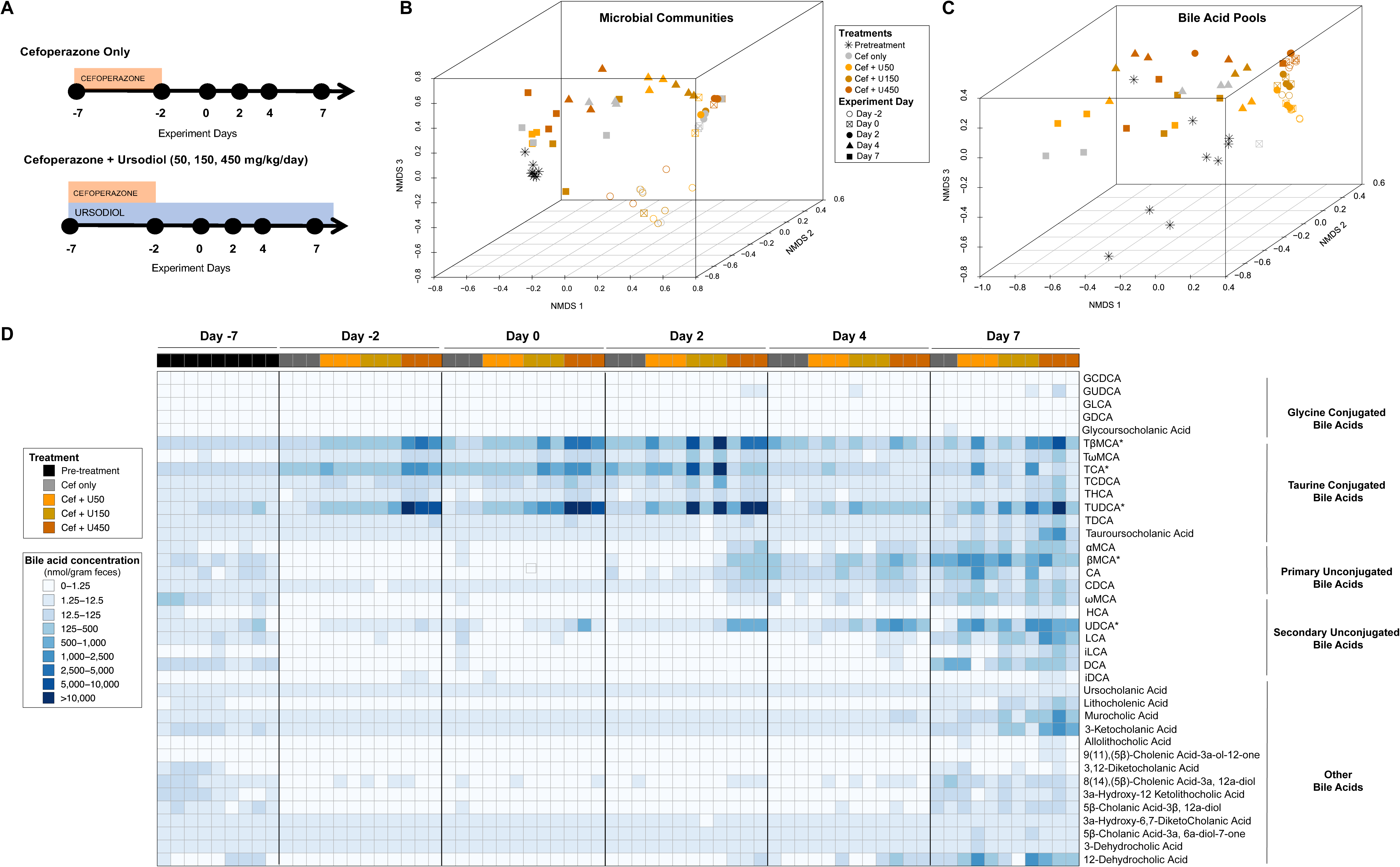
Ursodiol does not alter the gut microbiota but does alter the bile acid metabolome. Groups of C57BL/6J mice (n=8 mice/treatment group) were **(A)** given cefoperazone in their drinking water ad libitum for 5 days alone and or with ursodiol at three doses (50, 150, and 450 mg/kg) given daily via oral gavage for 14 days. An n = 3 mice/treatment group were selected for further microbiome and bile acid metabolomic analysis. **(B)** NMDS ordination of the fecal microbial communities from mice treated and untreated with ursodiol was calculated from Yue and Clayton dissimilarity metric (θ_YC_) on OTU at a 97% cutoff. **(C)** NMDS ordination illustrates dissimilarity indices via Horn distances between the bile acid profiles of fecal samples from treated and untreated mice. **(D)** A heatmap of bile acid concentrations present in nanomoles (nmol) per gram of feces, ranging from 0 to 38221.91 (nmol/g feces). Asterisks next to bile acids represent ones that were significantly altered during ursodiol treatment.

The fecal microbial community structures of all treatment groups were significantly different from pretreatment at all time points, based on an analysis of molecular variance (AMOVA) performed for each day (Fig. 2B). There were no significant differences between the fecal microbial community structures based on ursodiol pretreatment. The shifts in the fecal microbiota seen in the NMDS were predominantly due to the cefoperazone treatment and experiment day, and not as a result of the ursodiol pretreatment.

Targeted bile acid analysis via UPLC-MS/MS detected 38 bile acids in the feces of mice. During the first 9 days of the experiment (Day −2 to Day 7), the fecal bile acid profiles of mice cluster based on the dose of ursodiol administered (Fig. 2C). Ursodiol dramatically alters the fecal bile acid metabolome compared to pretreatment as seen in Fig. 2D and Table S3. Bile acids that significantly changed over the course of the experiment in mice that received ursodiol compared to mice that only received cefoperazone included TβMCA, TCA, TUDCA, βMCA, and UDCA (Fig. S2, two-way ANOVA with Tukey’s multiple comparisons *post hoc* preformed on each experiment day, p values ranged from p <0.0001 to p = 0.0012).

### Ursodiol attenuates disease in mice early during *C. difficile* infection

Due to potent inhibitory effect of ursodiol on *C. difficile in vitro*, and increased gastrointestinal bile acids *in vivo*, we sought to determine the extent to which pretreatment with ursodiol influences the natural course of CDI in a reproducible mouse model. Since supra-physiologic UDCA intestinal concentrations were achieved in Fig. 2D when mice were dosed with 450 mg/kg/day of ursodiol, we selected this dose.^58^ All C57BL/6J mice in each group received cefoperazone in their drinking water from Day −7 to Day −2 with a two day washout on normal drinking water. Control groups included two uninfected groups that received cefoperazone no ursodiol (Cef only), and cefoperazone with ursodiol (Cef + U450) (Fig. 3A). An additional control group received cefoperazone no ursodiol, with *C. difficile* challenge (*C. diff* only). Mice were orally gavaged with ursodiol (450 mg/kg in corn oil vehicle) daily for 7 days prior to challenge with *C. difficile* on Day 0, and throughout the infection (*C. diff* + U450) (Fig. 3A).

**Figure 3:**
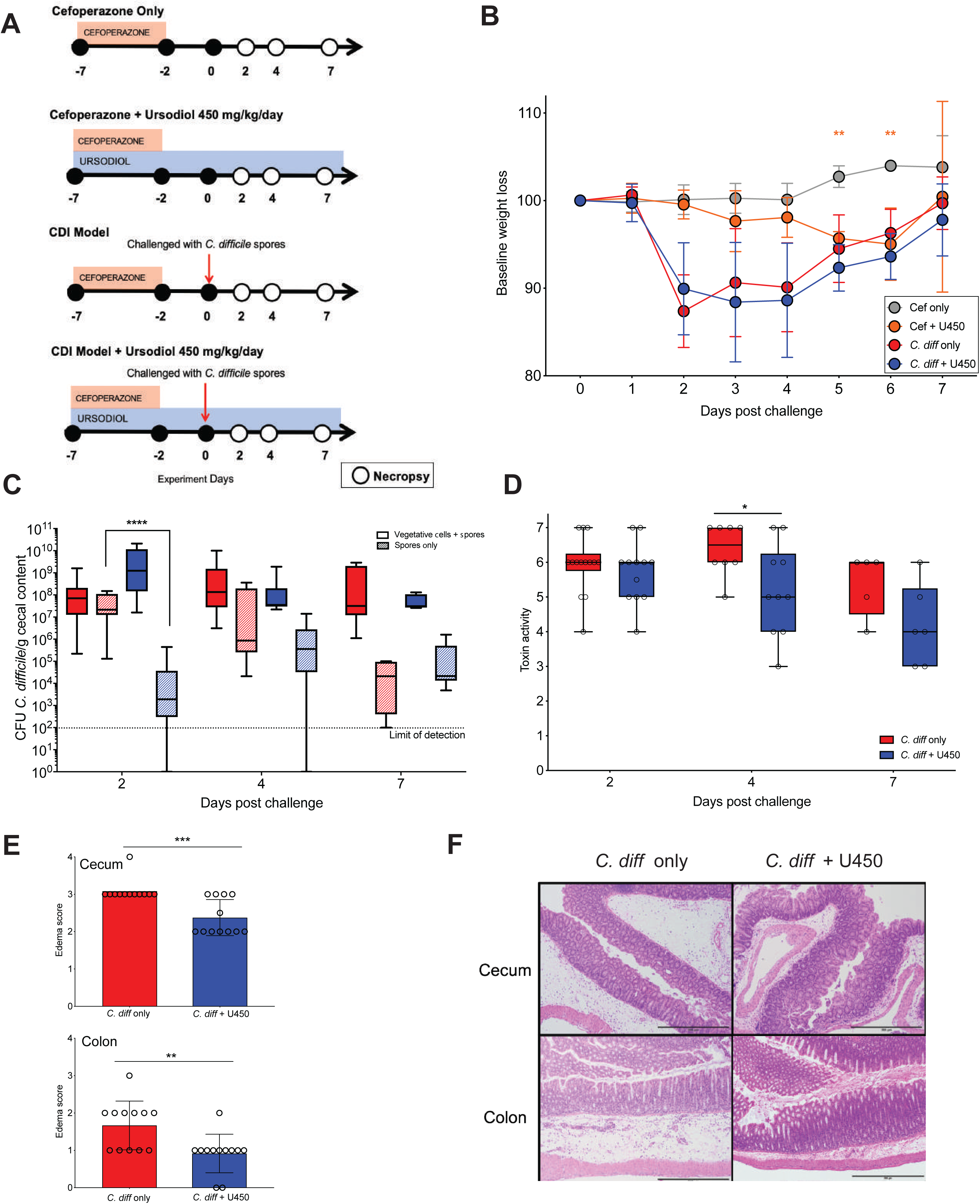
Ursodiol attenuates disease early during *C. difficile* infection. **(A)** Groups of C57BL/6J mice (n=12 mice/treatment group) were given cefoperazone in their drinking water *ad libitum* for 5 days alone and or with ursodiol at 450 mg/kg given daily via oral gavage for 14 days. Two groups of mice were then challenged with 10^5^ spores of *C. difficile* via oral gavage at day 0. Mice were monitored for weight loss and clinical signs of CDI from day 0 to day 7. Necropsy was performed on days 2, 4, 7 (open circles). **(B)** Baseline weight loss of mice challenged with and without *C. difficile*, and pretreated with and without ursodiol. Significance was determined by a two-way ANOVA with Sidak’s multiple comparisons *post hoc* test. **(C)** *C. difficile* bacterial load in the cecal content throughout CDI. Solid boxes represent total vegetative cells and spores and hashed boxes represent spores only. Significance was determined by a two-way ANOVA with Tukey’s multiple comparisons *post hoc* test. **(D)** Vero cell cytotoxicity assay from cecal content throughout CDI. Significance was determined by a Student’s t-test corrected with the Holm-Sidak method for multiple comparisons. **(E)** Histopathologic scoring of edema in the murine cecum and colon at day 2 post challenge. Significance was determined by a Student’s t-test corrected with the Holm-Sidak method for multiple comparisons. Error bars represent the standard deviations from the mean. **(F)** Representative histomicrograph images of H&E stained cecum (upper) and colon (lower) for groups indicated, shown at 100X total magnification with 500 µm scalebar. For all graphs (*, p ≤ 0.05; **, p ≤ 0.01; ***, p ≤ 0.001; ****, p ≤ 0.0001).

Two groups of mice (*C. diff* only and *C. diff* + U450) were challenged with 10^5^ spores of *C. difficile* R20291 on Day 0 and within 24 hr were colonized with greater than 10^7^ CFUs per gram of feces (data not shown). Mice challenged with *C. difficile* with and without ursodiol pretreatment lost a significant amount of their baseline weight by Day 2 post challenge (*C. diff* only, red circles and *C. diff* + U450, blue circles Fig. 3B). Additionally, there was no significant difference in weight loss between these groups over the course of the experiment (based on two-way ANOVA). However, mice pretreated with ursodiol (*C. diff* + U450) had a 24-36 hr delay in clinical signs (lethargy, inappetence, diarrhea, and hunched posture) related to CDI compared to untreated mice (*C. diff* only). Uninfected control mice (Cef only and Cef *+* U450) revealed no clinical signs of disease during the experiment and remained *C. difficile* free. The Cef only group of mice displayed weight gain over the course of the experiment as expected (Fig. 3B, gray circles). The Cef + U450 group did have significant weight loss compared to Cef only mice at Day 5 and Day 6 of the experiment (two-way ANOVA with Sidak’s multiple comparisons *post hoc* test; p = 0.0013, p = 0.0001 respectively; Fig. 3B, orange circles). However, no significant difference was noted between these groups by Day 7 (Fig. 3B). An additional control group of mice challenged with *C. difficile* and pretreated with the vehicle corn oil alone (*C. diff +* corn oil) displayed no significant difference in disease outcome compared to the *C. diff* only group (Fig. S3).

Mice pretreated with ursodiol (*C. diff* + U450) had equivalent total cecal bacterial loads of *C. difficile* (CFU/g cecal content) compared to untreated mice (*C. diff* only) (Fig. 3C). However, mice pretreated with ursodiol had a significant reduction in *C. difficile* spores within cecal content at Day 2 compared to untreated mice (log transferred CFU/g content, two-way ANOVA with Tukey’s multiple comparisons *post hoc* test, p < 0.0001; Fig. 3C). Mice ceca remained persistently colonized with *C. difficile* throughout the 7 day experiment. At Day 4, *C. difficile* toxin activity within cecal content was significantly reduced in ursodiol pretreated compared to untreated mice (Student’s t-test corrected with the Holm-Sidak method for multiple comparisons, p = 0.014; Fig. 3D).

Histopathologic evaluation of cecal and colonic tissue revealed no significant difference in total histologic score between ursodiol pretreated and untreated mice (Fig. S4). When assessing individual parameters, there was no significant difference between epithelial damage or inflammatory infiltrate between the treatment groups. At Day 2, mice pretreated with ursodiol had a significant reduction in edema score (scored 0-4) in both cecal and colonic tissue compared to untreated mice (Student’s t-test corrected with the Holm-Sidak method for multiple comparisons, p = 0.00025, p = 0.0049 respectively; Fig. 3E and F).

### Ursodiol alters the host inflammatory transcriptome early during CDI

Since there were minimal changes in the *C. difficile* life cycle early during infection to explain the decrease in edema, we wanted to examine the impact of ursodiol pretreatment on the host inflammatory response during CDI. We interrogated transcriptional changes related to the cecal and colonic inflammatory response at Day 2 post challenge using a murine specific panel of inflammatory markers (Nanostring nCounter) that included 264 transcripts. After processing the raw data, mRNA log2 normalized z-scores for 261 genes were obtained and genes significantly altered by ursodiol pretreatment compared to untreated mice with CDI were evaluated. Ursodiol mediated alterations in gene expression fold change were different between cecal and colonic tissue (Fig. 4 and Fig. S5). In cecal tissue, 173 genes had a significant expression fold change when comparing ursodiol pretreated mice to untreated mice, with increased expression in 74 genes and decreased expression in 99 genes (Fig. 4A and B, Table S4 and S5). In colonic tissue, 139 genes had a significant expression fold change when comparing ursodiol pretreated to untreated mice, with increased expression in 58 genes and decreased expression in 81 genes (Fig. S5A and B, Table S4 and S5).

**Figure 4:**
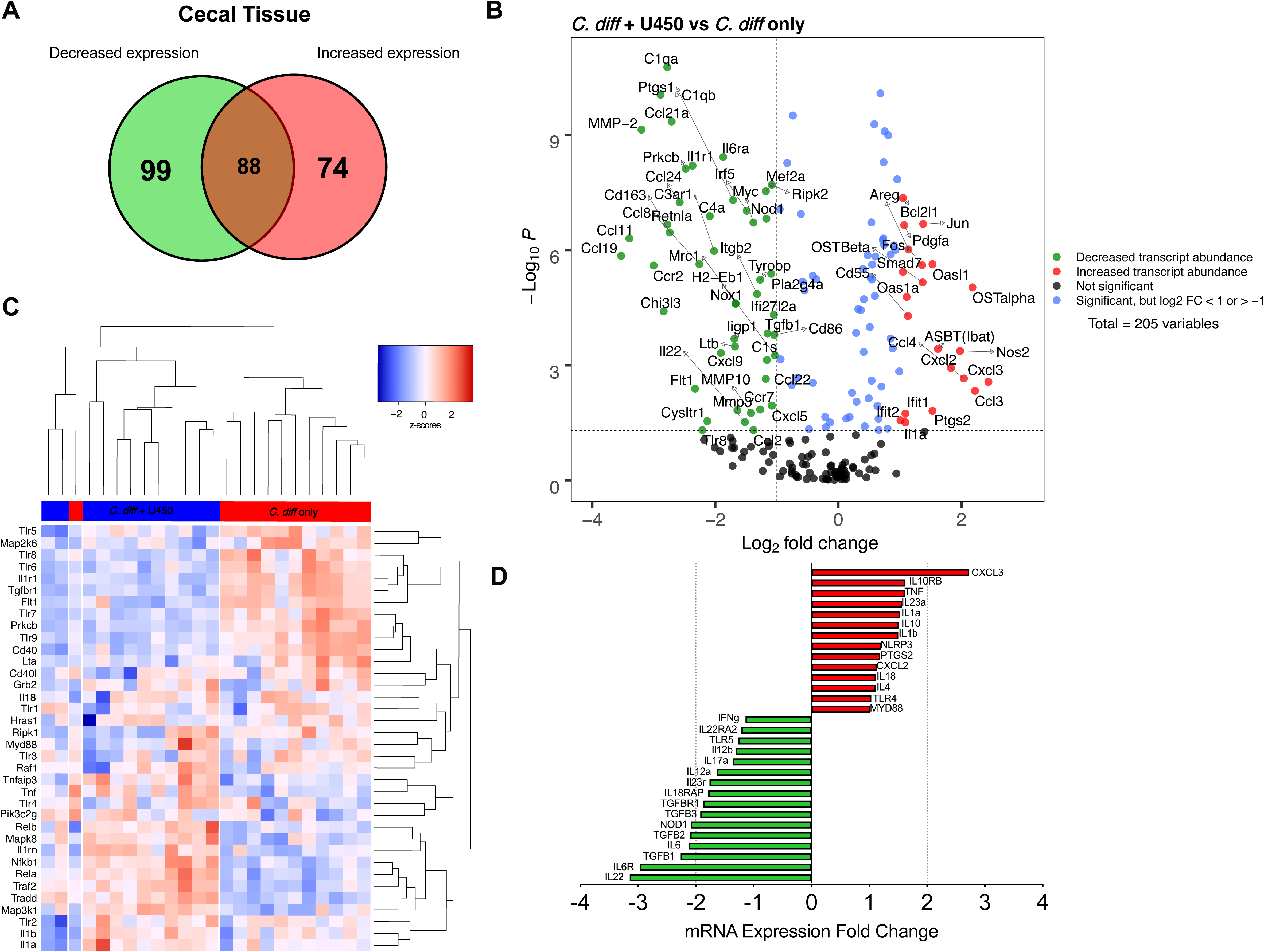
Ursodiol alters the host inflammatory transcriptome of the cecum early during CDI. **(A)** Venn diagram depicting the mRNA expression fold change comparing the ursodiol pretreated compared to untreated mice in cecal tissue at day 2 post challenge with *C. difficile*, from two independent experiments performed with a total of n=12 mice/treatment group. Of the 261 genes evaluated, a total of 173 cecal genes had significant gene expression fold changes (increased expression, red; decreased expression, green). **(B)** Volcano plots highlighting genes whose transcript levels changed by greater than 1-fold and met the significant threshold *p* ≤ 0.05. Genes highlighted in red had increased transcript levels, while those highlighted in green had decreased levels. Blue points represent genes whose results were significant but did not met the specify fold change. Black points represent genes whose results failed to meet the significance threshold. **(C)** Cecal inflammatory transcriptome heatmap of calculated z-scores for log2 transformed count data of mRNA expression levels (y-axis) comparing ursodiol pretreated (blue) to untreated mice (red). Dendrograms represent unsupervised clustering. **(D)** mRNA expression fold differences in increased expression (red) and decreased expression (green) of genes involved with NF-kB signaling in cecal tissue comparing ursodiol pretreated to untreated mice.

Ingenuity Pathway Analysis (IPA) identified NF-κB signaling as an inflammatory canonical pathway in both cecum and colon significantly altered in ursodiol pretreated mice (p < 0.0001 for both tissues). Based on unsupervised clustering, genes involved with NF-κB signaling in ursodiol pretreated mice clustered distinctly and separately from untreated mice in both cecal and colonic tissue (Fig. 4C and Fig. S5C). Differences in individual gene expression fold changes were noted in NF-κB signaling between the cecal and colonic tissue (Fig. 4D and Fig. S5D). In both tissues, components important for mediating the host inflammatory response to CDI, such as IL-1R1 and various TLRs, were significantly decreased in expression in the ursodiol pretreated mice compared to untreated mice at Day 2 (Fig. 4D and Fig. S5D, see Table S6 for p-values). Collectively, these results suggest that ursodiol pretreatment in CDI mice alters the inflammatory transcriptome in cecal and colonic tissue by impacting essential host inflammatory signaling pathways, such as NF-κB, early in the course of CDI.

### Ursodiol induces bile acid transport and signaling via FXR/FGF15 pathway

Since ursodiol treatment alters the bile acid metabolome in mice (Fig. 2D) and bile acids are known to modulate signaling via the nuclear bile acid receptor FXR and the plasma membrane receptor TGR5^59^, we sought to define transcriptional changes in bile acid transport and signaling via the FXR/FGF15 pathway. We selected seven genes in the FXR/FGF15 pathway to interrogate within cecal and colonic tissue using a customized murine panel (Nanostring nCounter). Bile acids within the gastrointestinal tract are transported back to the liver via enterohepatic recirculation by several active transporters, ASBT on the apical surface, and OSTα/β on the basolateral surface of enterocytes (Fig. 5C)^59^. Cytoplasmic Ibabp helps to facilitate bile acid transport (Fig. 5C)^59^. Bile acids in ileal enterocytes and colonic L cells bind to FXR and activate the FXR-RXR heterodimer complex, resulting in transcription of target genes, such as FGF15/19. FGF15/19 is secreted into the portal vein and transported back to the liver, where it inhibits expression of *CYP7A1* in hepatocytes to directly regulate bile acid synthesis.^59^

**Figure 5:**
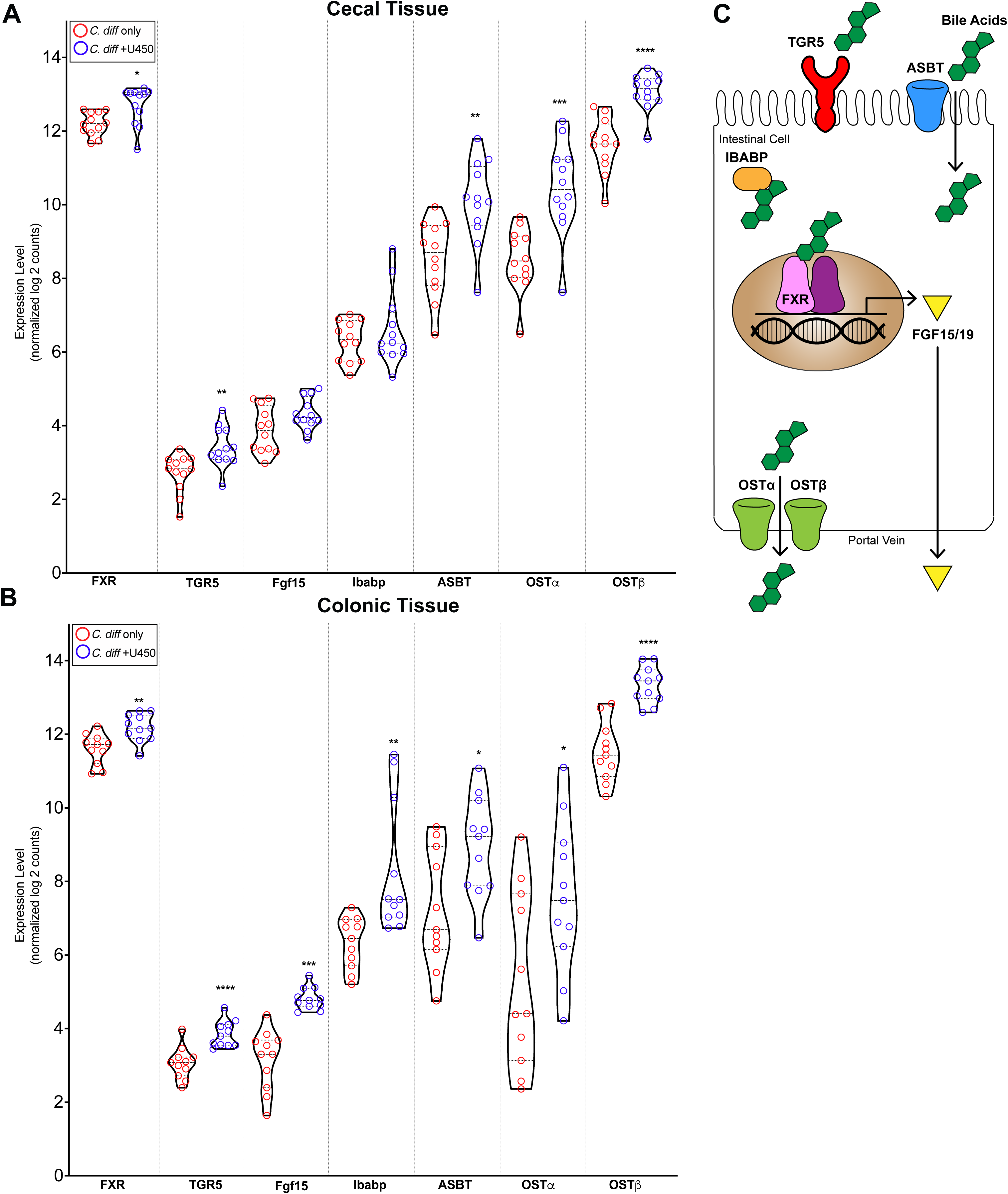
Ursodiol alters the FXF-FGF15 transcriptome during CDI. Violin plots of **(A)** cecal and **(B)** colonic mRNA expression levels of normalized log2 counts of seven genes involved with the FXF-FGF15 pathway. Significance determined by a Student’s parametric t-test with Welch’s corrected (*, p ≤ 0.05; **, p ≤ 0.01; ***, p ≤ 0.001; ****, p ≤ 0.0001). **(C)** Schematic of seven genes within the FXF-FGF15 pathway. Bile acids modulate signaling via the nuclear bile acid receptor FXR (farnesoid X receptor) and plasma membrane receptor TGR5 (or G-protein coupled bile acid receptor Gpbar1). Bile acid within the gastrointestinal tract are transported back to the liver via enterohepatic recirculation by several active transporters, ASBT (apical sodium dependent bile acid transporter) on the apical surface, and OSTα/β (organic solute transporters) on the basolateral surface of enterocytes. Cytoplasmic Ibabp (ileal bile acid-binding protein) helps to facilitate bile acid transport. Bile acids in ileal enterocytes and colonic L cells bind to FXR and activate the FXR-RXR heterodimer complex, resulting in transcription of target genes, such as FGF15/19. FGF 15/19 is secreted into the portal vein and transported back to the liver, where it inhibits expression of *CYP7A1* in hepatocytes to directly regulate bile acid synthesis.

In cecal tissue, mRNA expression of the receptors FXR and TGR5, and the bile acid transporters ASBT and OSTα/β were significantly increased in ursodiol pretreated mice compared to untreated mice at Day 2 post challenge (Fig. 5A, Student’s parametric t-test with Welch’s correction, see Table S7 for p values). In colonic tissue, mRNA expression of the receptors FXR and TGR5, the signaling molecule FGF15, and the bile acid transporters ASBT and OSTα/β were significantly increased in ursodiol pretreated mice compared to untreated mice (Fig. 5B, Student’s parametric t-test with Welch’s correction, see Table S7 for p values). Collectively, these results demonstrate that pretreatment of ursodiol in mice with CDI increases the expression of the FXR/FGF15 pathway transcriptome in both cecal and colonic tissue.

## Discussion

Ursodiol, the FDA approved commercial formulation of UDCA, is used to treat a variety of hepatic and biliary diseases including: cholesterol gallstones, primary biliary cirrhosis, primary sclerosing cirrrhosis, cholangitis, non-alcoholic fatty liver disease, chronic viral hepatitis C, and cholestasis of pregnancy.^31–40^ However, limited evidence is available for use of ursodiol in CDI patients. To date, only a single case report demonstrating successful prevention of recurrent *C. difficile* ileal pouchitis with oral UDCA administration is available,^29^ suggesting that exogenous administration of the secondary bile acid UDCA may be able to restore colonization resistance against *C. difficile in vivo*.^29^ Although the mechanisms by which ursodiol exerts its impact on *C. difficile* are unknown, off-label use of ursodiol is currently in Phase 4 clinical trials for the prevention of recurrent CDI.^41^ In the present study, ursodiol pretreatment impacted the life cycle of *C. difficile in vivo*. Although, *C. difficile* bacterial load and toxin activity was not affected at day 2, pretreatment with ursodiol resulted in a 24-36 hr delay in the clinical course of CDI, suggesting attenuation of CDI pathogenesis early during infection. Although ursodiol pretreatment did not alter the gut microbiota composition, significant modifications in the bile acid metabolome were observed. Several key bile acid species were significantly increased following ursodiol administration, namely UDCA, TUDCA, and TβMCA. The tissue inflammatory transcriptome was altered, including gene expression of bile acid activated receptors nuclear farnesoid X receptor (FXR) and transmembrane G protein-coupled membrane receptor 5 (TGR5), which are able to modulate the innate immune response through signaling pathways such as nuclear factor *κ*-B (NF-κB). This is the first study to investigate the impact of innate immune response regulation by bile acid activated receptors, FXR and/or TGR5, during CDI.

Modulation of bile acid pools to restore colonization resistance against *C. difficile* is a rational therapeutic option, based on the importance bile acids play in the life cycle of *C. difficile.*^25, 27^ Traditional therapeutic approaches to CDI have focused on restoring bile acid pools by reestablishing the entire microbial community, such as with fecal microbiota transplantation.^13, 28^ Or supplementation of key commensals, such as *Clostridium scindens*, a bacterium capable of synthesizing microbial derived secondary bile acids, which when supplemented leads to partial restoration of colonization resistance against *C. difficile*.^11^ In the present study, we deduced that direct supplementation of bile acids, such as UDCA or ursodiol which are known to inhibit the life cycle of *C. difficile,*^26^ may be sufficient to restore colonization resistance against this enteric pathogen by directly impacting *C. difficile,* but also by modulating the host inflammatory response.

The host innate immune response to CDI is an essential component of *C. difficile* pathogenesis.^1^ The acute host innate immune response is initiated by *C. difficile* toxin-mediated damage to the intestinal epithelial cells, loss of the epithelial barrier, and subsequent translocation of luminal gut microbes.^1, 60^ *C. difficile* toxin A (TcdA) and toxin B (TcdB), directly disrupt the cytoskeleton of intestinal epithelial cells, resulting in disassociation of tight junctions and loss of epithelial integrity.^1, 61^ In response to toxin-mediated destruction, resident immune cells and intoxicated epithelial cells release pro-inflammatory cytokines and chemokines to recruited circulating innate immune cells and adaptive immune cells (reviewed in Abt et al.).^1^ A prominent inflammatory pathway that is activated in intestinal epithelial cells by TcdA and TcdB is the NF*κ*-B signaling pathway through the phosphorylation of mitogen-activated protein kinase (MAPK), which leads to the transcription of pro-inflammatory cytokines, including IFN-*γ*, TNF-*α*, IL-6, and IL-1*β*.^62–65^ In the present study, pretreatment with ursodiol resulted in alterations in the host intestinal inflammatory transcriptome. Genes involved with NF-*κ*B signaling pathway in both cecum and colon, segregated based on ursodiol pretreatment. This is not surprising given the broad ursodiol mediated alterations in bile acid metabolome observed and the building evidence that bile acid activated receptors regulate the host innate immune response (recently reviewed in Fiorucci et al. 2018).^44^ In particular, interactions between the gut microbiota and bile acids result in alterations in bile acid pools, which in turn modulate signaling via the FXR and TGR5, both of which can regulate the innate immune response.^44, 59, 66^ Bile acid activation of TGR5 involves cAMP-protein kinase A (PKA)-mediated inhibition of NF-*κ*B, and bile acid activation of FXR leads to FXR-nuclear receptor corepressor 1 (NCor1)-mediated repression of NF-*κ*B responsive elements (NRE). Bile acid activation of both pathways blunt NF-*κ*B gene transcription including pro-inflammatory cytokines IFN-*γ*, TNF-*α*, IL-6, and IL-1*β*.^67, 68^ Additionally, TGR5 directly regulates IL-10 expression by a cAMP/PKA/p CAMP-responsive element binding protein pathway, thus further attenuating the inflammatory response.^69^

Bile acids differ in their affinity and agonistic/antagonistic effects for FXR and TGR5.^44^ We observed global alterations in bile acids that are both FXR and TGR5 agonists and antagonists, lending to the complexity of unravelling the cumulative effects on bile acid activated receptors and how this impacts the overall innate immune response during CDI. Based on FXR and TGR5 signaling transcriptomics, ursodiol pretreatment in the present study resulted in increased cecal and colonic expression of genes in these pathways during CDI. Collectively, this suggests that ursodiol induced alterations in the bile acid metabolome results in alterations of host inflammatory transcriptome, including alterations in NF-*κ*B signaling pathway, likely via the bile acid activated receptors FXR and TGR5. Furthermore, this principle has also recently been demonstrated in recurrent CDI patients receiving FMTs, where effectiveness of FMT was associated with an upregulation of the FXR-FGF pathway.^70^

Whilst the present study demonstrated ursodiol-mediated attenuation of CDI early in the disease course, complete restoration of colonization resistance against this enteric pathogen was not achieved. Several limitations of the use of ursodiol pretreatment prior to CDI need to be considered. First, once daily dosing may be sufficient to obtain supraphysiologic pulses of UDCA, but these concentrations may not be sustained in the gastrointestinal tract throughout the day. Consequently, *C. difficile in vivo* is exposed to a daily pulse of ursodiol that inhibits its life cycle, similar to our *in vitro* results, only during a limited window. Alterations in the dosing scheme such as dividing the dose over two or four administration times may ameliorate this potential limitation. Secondly, alterations in the inflammatory microenvironment during the initial phases of CDI were observed in this study, however these effects were not sustained throughout CDI. Potentially, longer pretreatment with ursodiol is needed to result in sustained alterations to the host innate immune response. Lastly, ursodiol may be more effective as a preventative for relapsing CDI, meaning that following successful treatment of CDI with antibiotic treatment ursodiol could be administered to prevent relapse of infection.

Regardless, our results highlight that ursodiol pretreatment was able to significantly alter the bile acid metabolome and host inflammatory transcriptome during CDI. Although ursodiol pretreatment did not result in colonization resistance against *C. difficile*, attenuated pathogenesis of *C. difficile* was noted early in the disease course. Collectively, we theorize that ursodiol induced alterations within the intestinal bile acid metabolome result in activation of bile acid receptors, such as FXR and TGR5, which are able to modulate the innate immune response through signaling pathways such as NF-*κ*B, thus mitigating an overly robust host inflammatory response that can be detrimental to the host. Given these results and the effectiveness of ursodiol to inhibit the *C. difficile* life cycle, ursodiol remains a viable non-antibiotic treatment and/or prevention strategy against CDI. Moreover, targeting bile acid activated receptors, FXR and TGR5, to attenuate an overly robust detrimental host immune response during CDI merits further consideration as a potential therapeutic intervention for CDI patients.

## Acknowledgements.

JAW was funded by the Ruth L. Kirschstein National Research Service Award Research Training grant T32OD011130 by NIH. This research was supported by work performed by The University of Michigan Microbial Systems Molecular Biology Laboratory (microbiome sequencing). The Microscopy Services Laboratory, Department of Pathology and Laboratory Medicine, is supported in part by P30 CA016086 Cancer Center Core Support Grant to the UNC Lineberger Comprehensive Cancer Center. CMT is funded by the National Institute of General Medical Sciences of the National Institutes of Health under award number R35GM119438. This project was also funded by an intramural grant from the North Carolina State University College of Veterinary Medicine.

## Figure legends

**Supplemental Figure 1: Effect of ursodiol on viability and spore formation of *C. difficile*.** Culture aliquots (100 μL) of *C. difficile* R20291, grown in BHI media, were taken over a 24 hr period and enumerated on TBHI plates to obtain total CFU/ml of total vegetative cells and spores (solid bars) and spores only (hashed bars). The positive controls (EthOH, gray bars) represent *C. difficile* grown in BHI media with ethanol and the treatments groups (blue bars) represents *C. difficile* grown in BHI media with varying concentrations of ursodiol from lowest to highest concentration (left to right). The limit of detection for this assay is 10^2^ CFU/ml. The data presented represents triplicate experiments. Significance was determined by Welch’s t-test using log transformed CFU/ml.

**Supplemental Figure 2: Ursodiol treatment alters the bile acid metabolome.** Box and whisker plots of bile acids that were significantly altered over the course of ursodiol treatment compared to mice that only received cefoperazone (based on a Two-way ANOVA with Tukey’s multiple comparisons post hoc test). An n=3 mice/treatment group were then selected for bile acid metabolomic analysis.

**Supplemental Figure 3: No effect with corn oil vehicle alone on CDI in mice. (A)** Baseline weight loss of cefoperazone treated mice challenged with *C. difficile* pretreated with (white, *C. diff* + corn oil) and without corn oil vehicle (red, *C. diff* only) (n=8 mice/treatment group). **(B)** *C. difficile* bacterial load in feces and **(C)** cecal content over the first 2 days post challenge. Solid boxes represent total vegetative cells and spores and hashed boxes represent spores only. Significance was determined by a two-way ANOVA test.

**Supplemental Figure 4: Total histologic scores were not significantly different with ursodiol pretreatment.** Histopathologic scoring of murine cecum and colon throughout CDI. Total histologic scores were calculated by adding all three scores from parameters assessed: epithelial damage, inflammation, and edema. Significance was determined by a two-way ANOVA with Sidak’s multiple comparisons *post hoc* test. Error bars represent the standard deviations from the mean.

**Supplemental Figure 5: Ursodiol alters the host inflammatory transcriptome of the colon early during CDI. (A)** Venn diagram depicting the mRNA expression fold change comparing the ursodiol pretreated to untreated mice in colonic tissue at day 2 post challenge with *C. difficile,* from two independent experiments performed with a total of n=12 mice/treatment group. Of the 261 genes evaluated, a total of 139 colonic genes had significant gene expression fold changes (increased expression, red; decreased expression, green). **(B)** Volcano plots highlighting genes whose transcript levels changed by greater than 1-fold and met the significant threshold *p* ≤ 0.05. Genes highlighted in red had increased transcript levels, while those highlighted in green had decreased levels. Blue points represent genes whose results were significant but did not met the specify fold change. Black points represent genes whose results failed to meet the significance threshold. **(C)** Colonic inflammatory transcriptome heatmap of calculated z-scores for log2 transformed count data of mRNA expression levels (y-axis) comparing ursodiol pretreated (blue) to untreated mice (red). Dendrograms represent unsupervised clustering. **(D)** mRNA expression fold differences in increased expression (red) and decreased expression (green) of genes involved with NF-kB signaling in colonic tissue comparing ursodiol pretreated to untreated mice.

**Supplemental Table 3:** Targeted bile acid metabolomics data from mice treated with cefoperazone with and without different doses of ursodiol.

**Supplemental Table 4:** Nanostrings differential gene expression data from cecal and colonic tissue comparing ursodiol pretreated to untreated mice.

**Supplemental Table 5:** Nanostrings z-score data from the cecal and colonic tissue comparing ursodiol pretreated to untreated mice.

**Supplemental Table 6:** Cecal and Colonic Tissue mRNA Expression Fold Change for NF-kB Signaling

**Supplemental Table 7:** Cecal and Colonic Tissue mRNA Expression Fold Change for FXR-FGF15 Pathway

---

**Supplemental Table 1:**
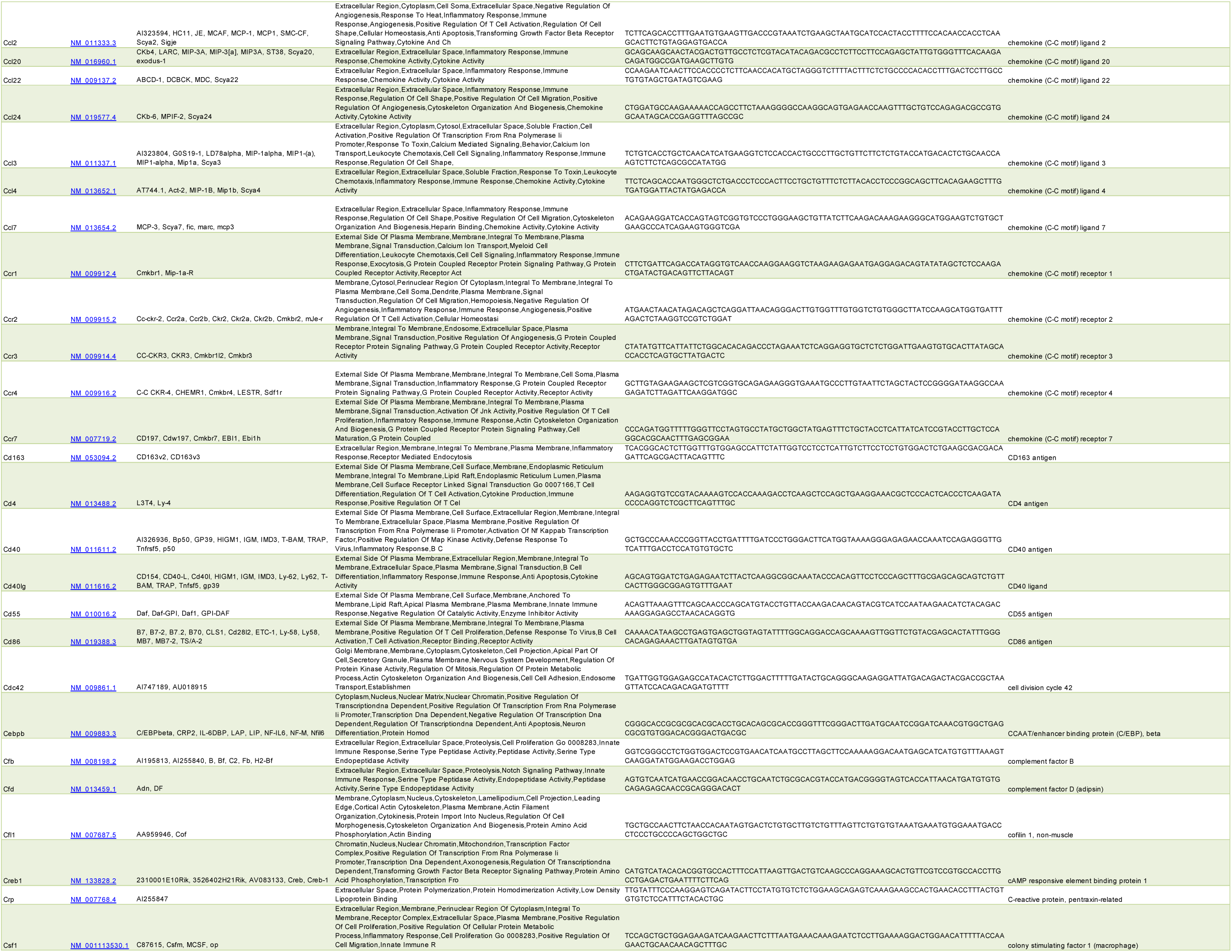
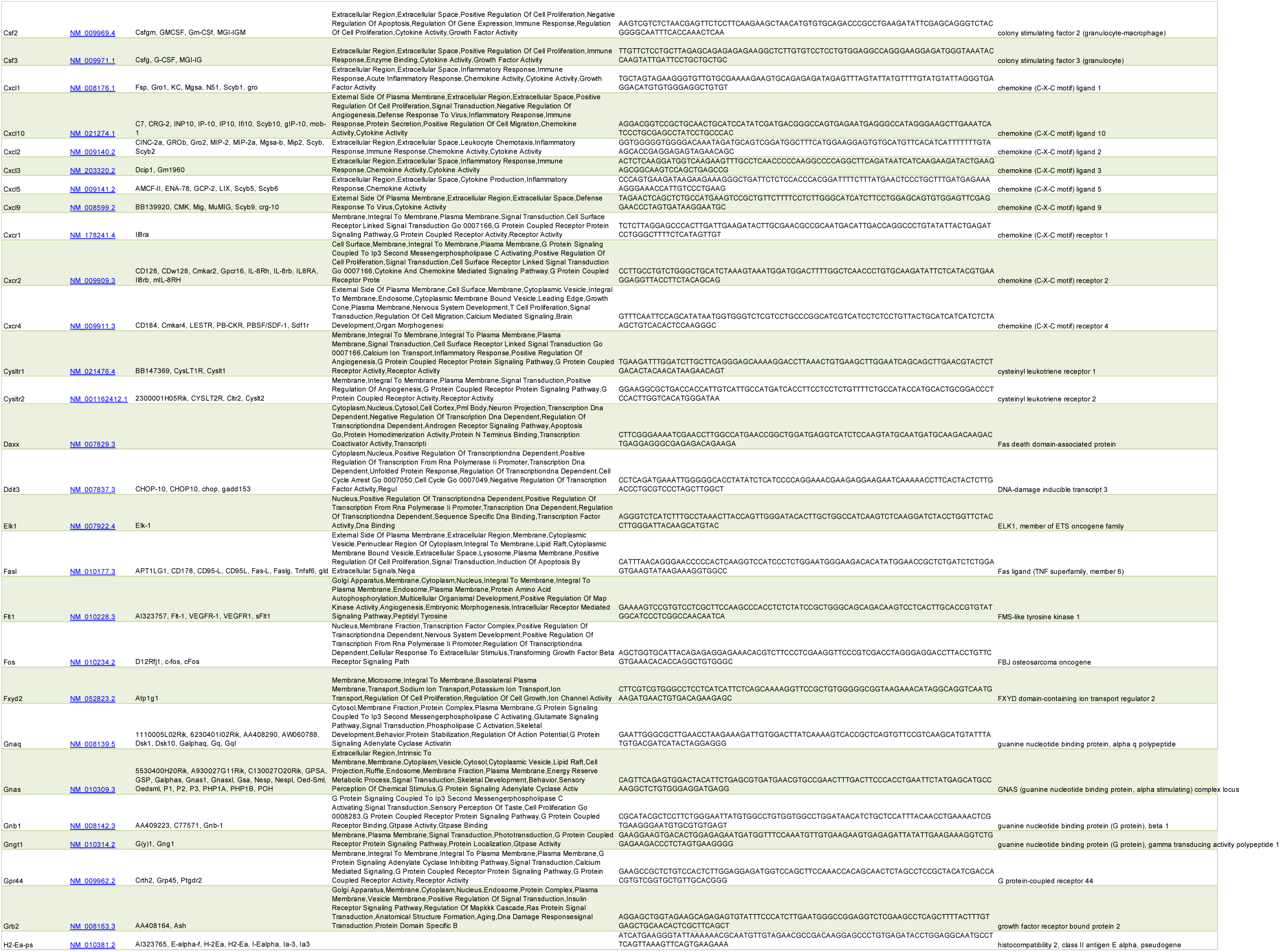
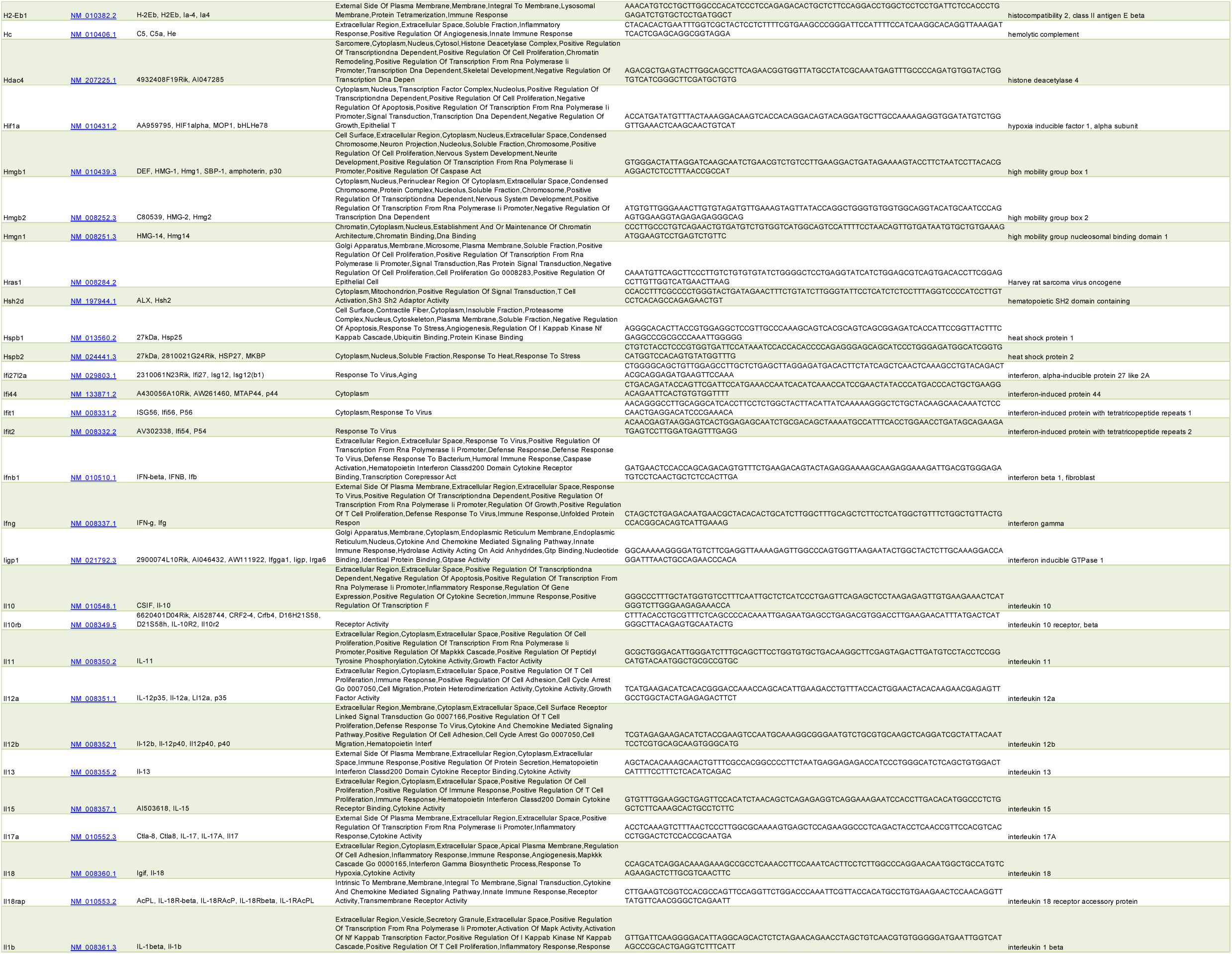
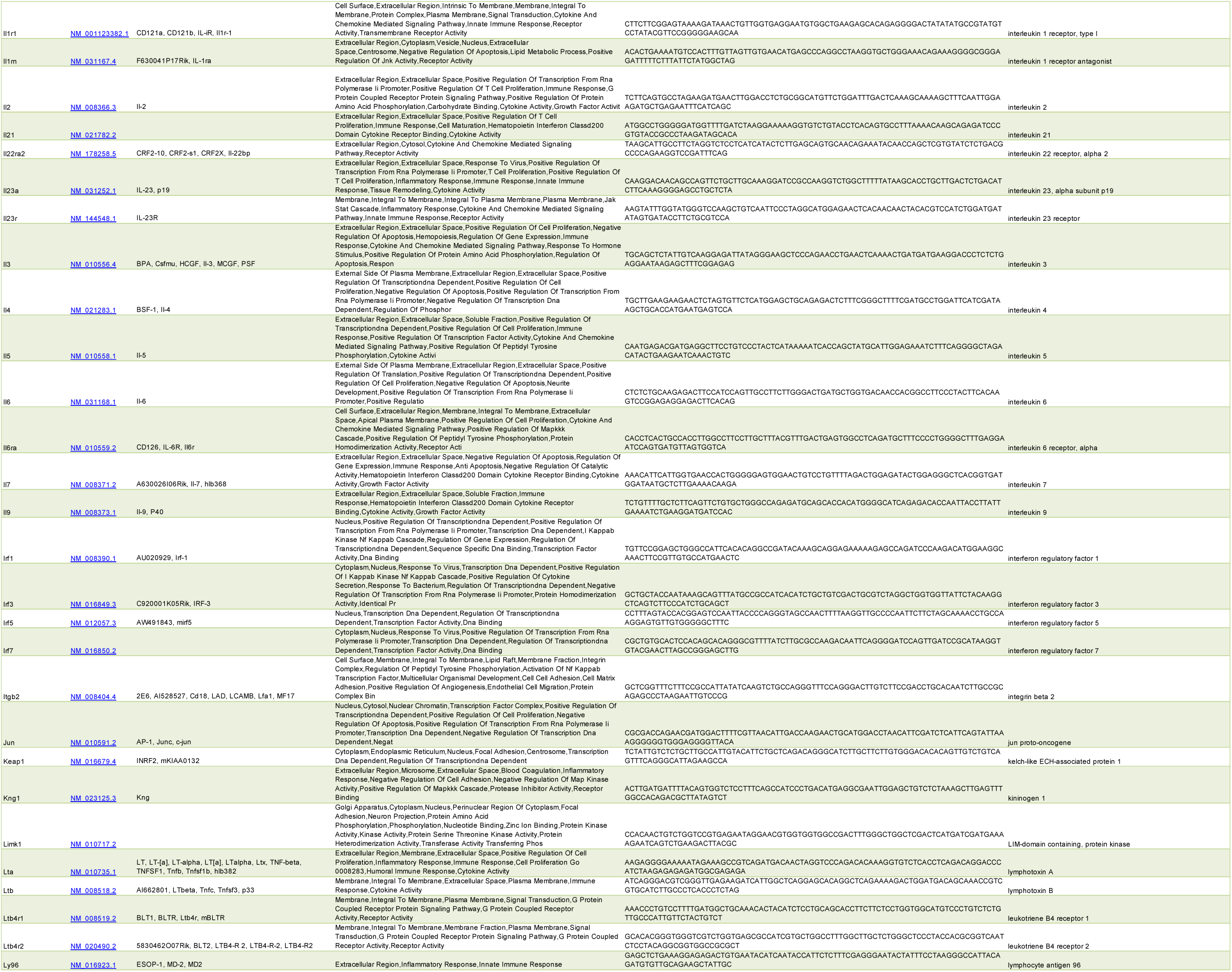
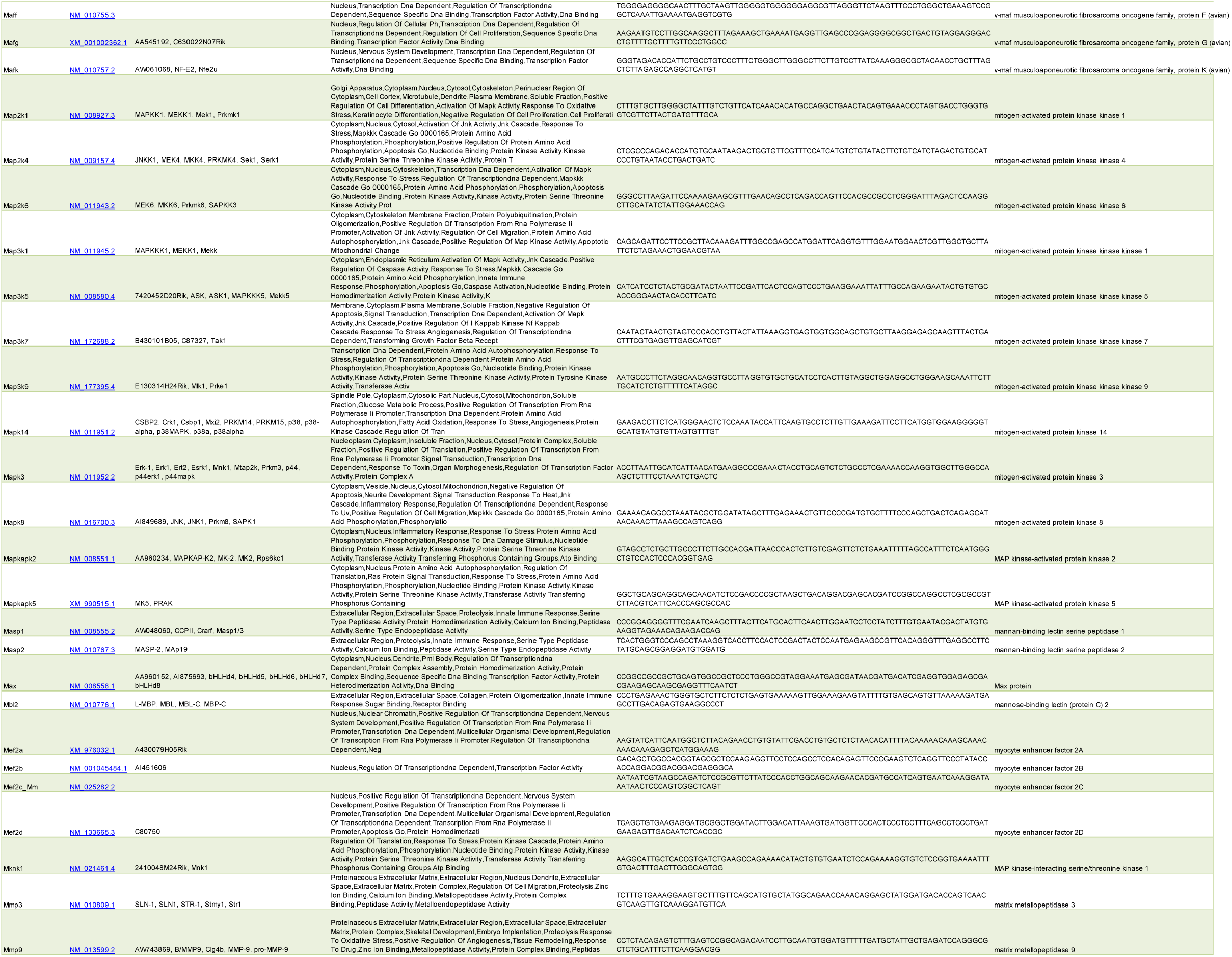
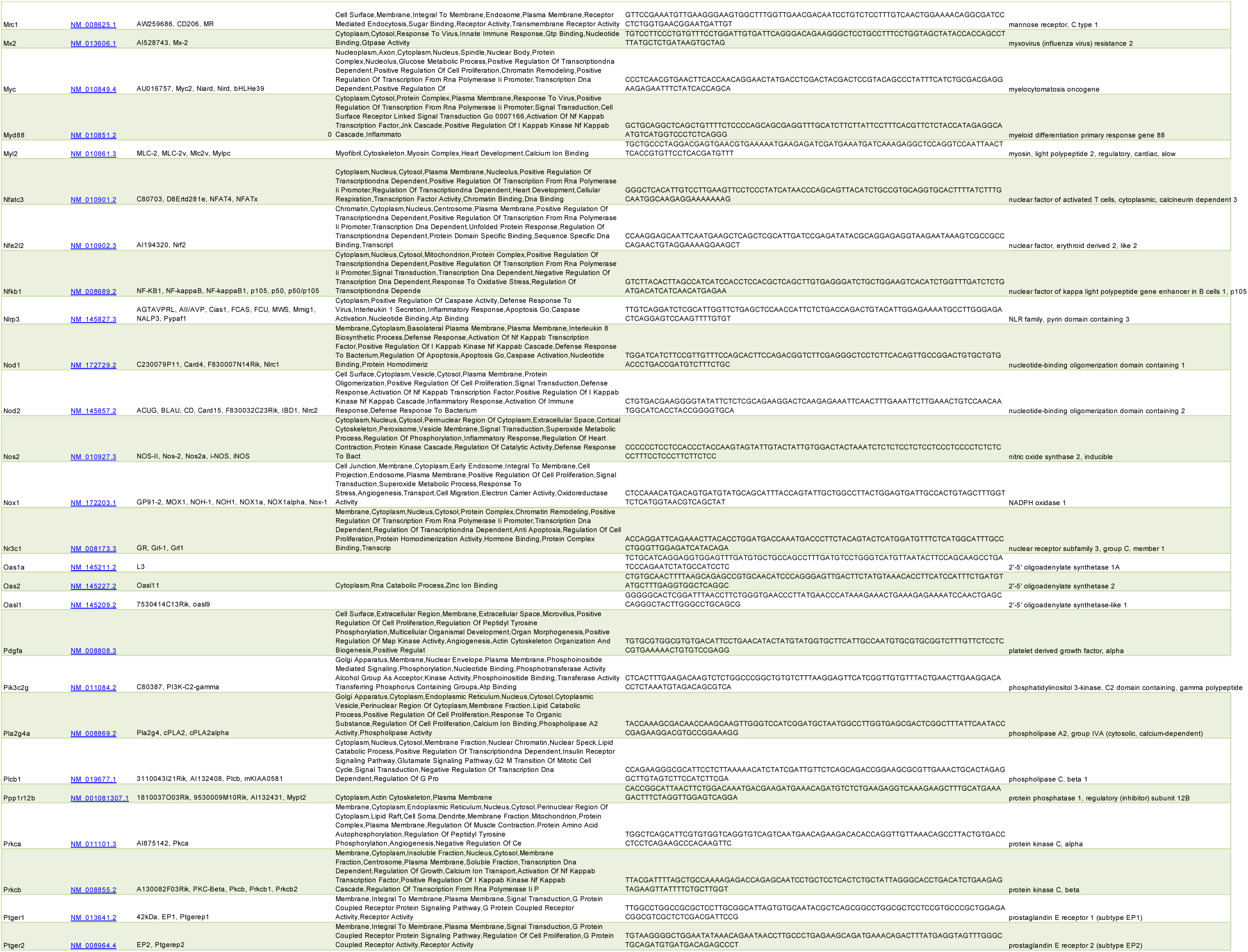
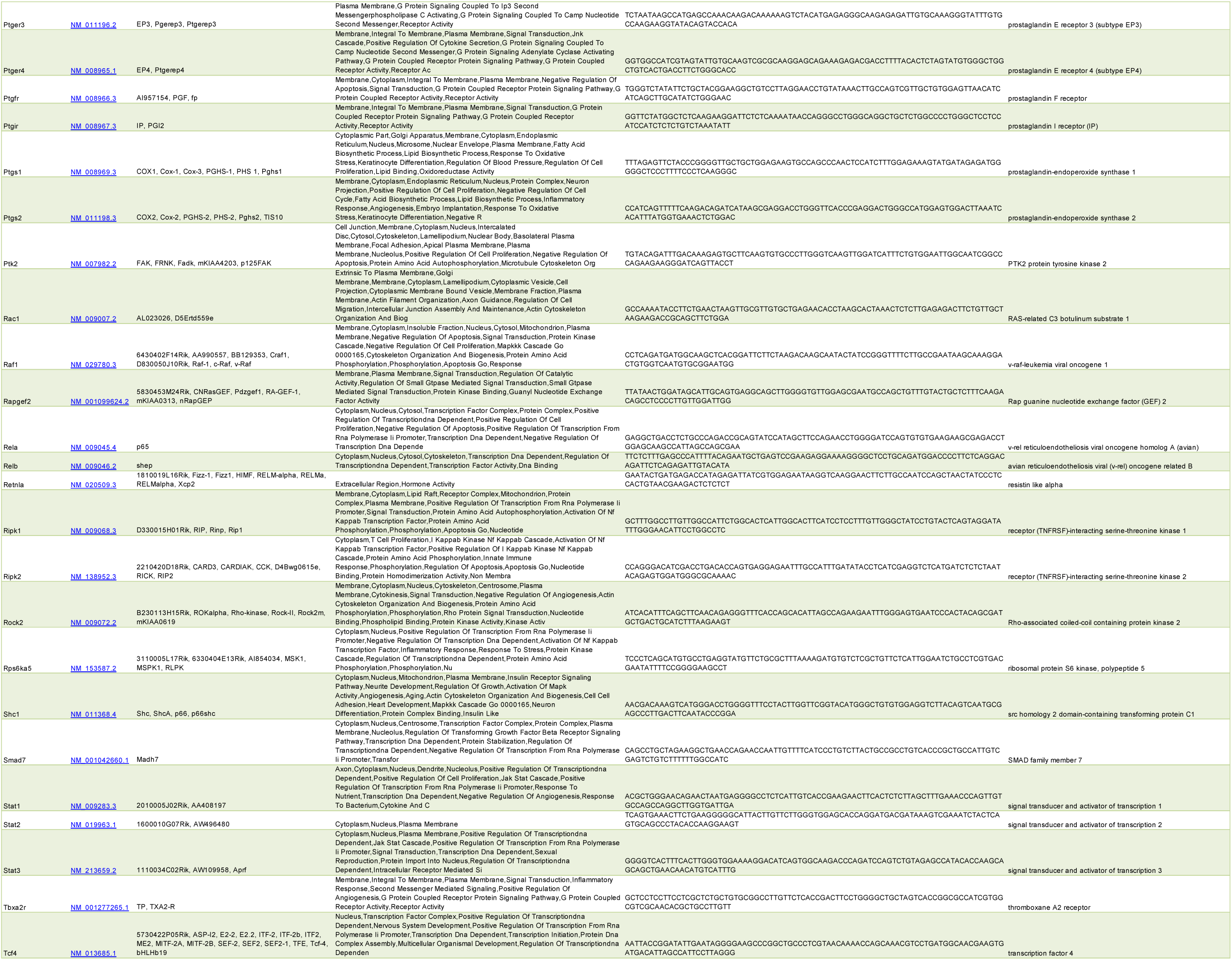
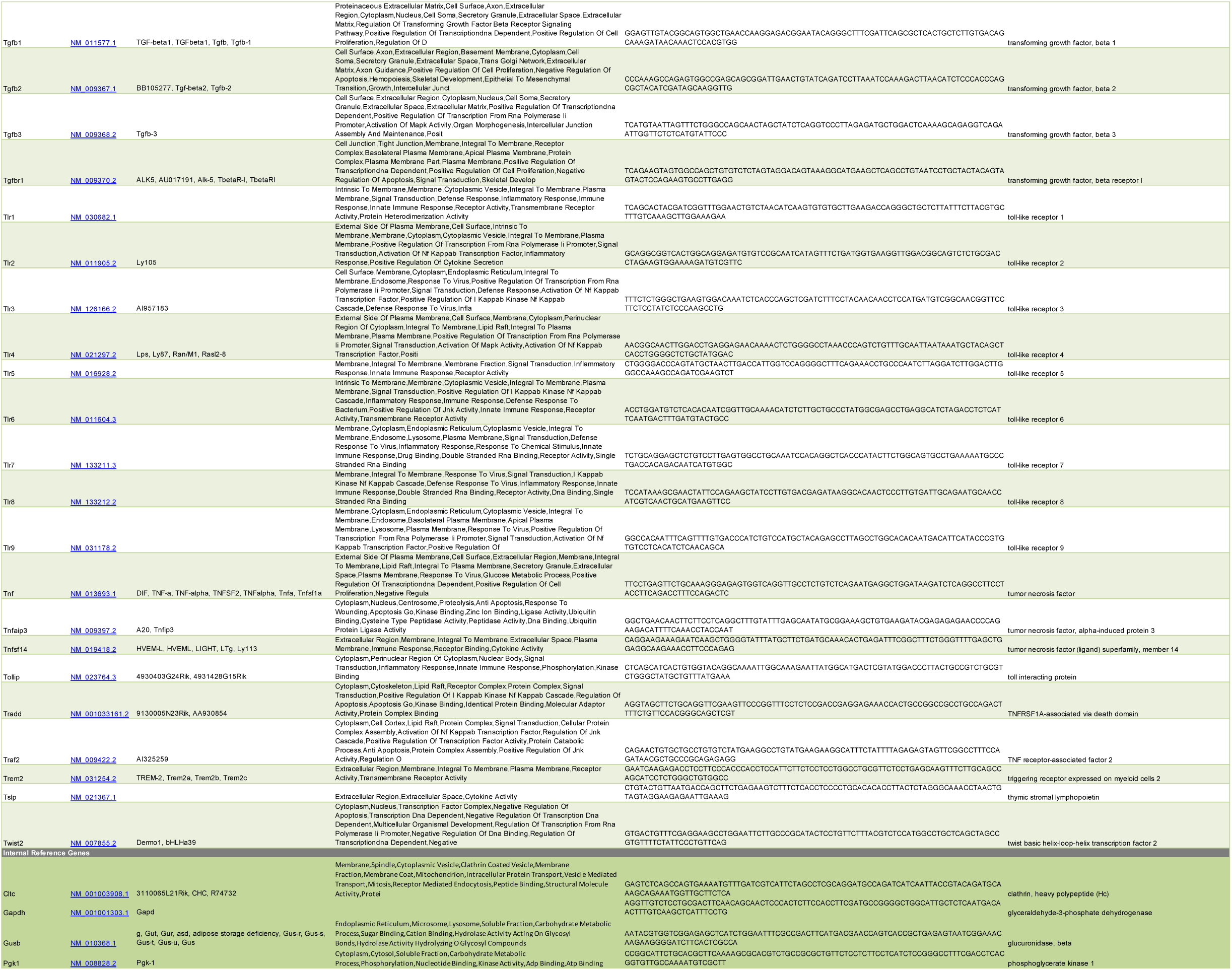
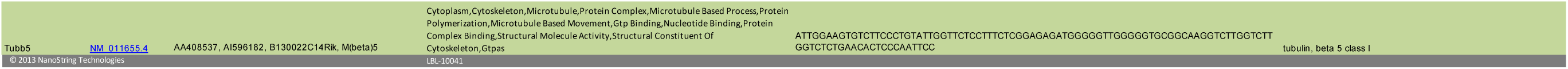
Nanostrings nCounter Mouse Inflammation V2 Panel Gene List (in word document)

**Supplemental Table 2:**
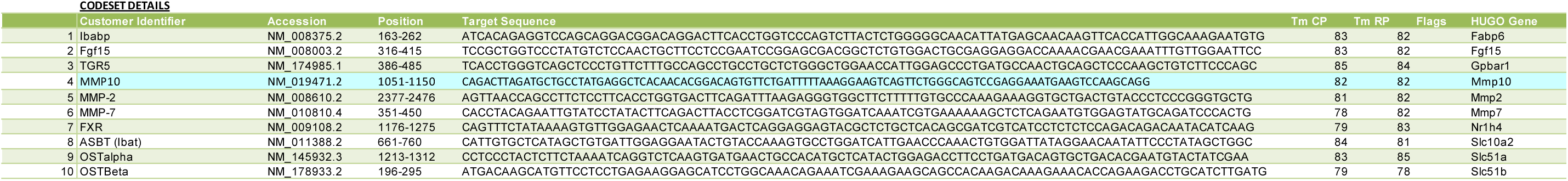
Nanostrings nCounter Mouse Custom Code Set Gene List (in word document)

